# RNA-Seq Analysis Reveals Localization-Associated Alternative Splicing across 13 Cell Lines

**DOI:** 10.1101/860783

**Authors:** Chao Zeng, Michiaki Hamada

## Abstract

Alternative splicing, a ubiquitous phenomenon in eukaryotes, is a regulatory mechanism for the biological diversity of individual genes. Most studies have focused on the effects of alternative splicing for protein synthesis. However, the transcriptome-wide influence of alternative splicing on RNA subcellular localization has rarely been studied. By analyzing RNA-seq data obtained from subcellular fractions across 13 human cell lines, we identified 8720 switching genes between the cytoplasm and the nucleus. Consistent with previous reports, intron retention was observed to be enriched in the nuclear transcript variants. Interestingly, we found that short and structurally stable introns were positively correlated with nuclear localization. Motif analysis reveals that fourteen RNA-binding protein (RBPs) are prone to be preferentially bound with such introns. To our knowledge, this is the first transcriptome-wide study to analyze and evaluate the effect of alternative splicing on RNA subcellular localization. Our findings reveal that alternative splicing plays a promising role in regulating RNA subcellular localization.

## 1 Introduction

Most eukaryotic genes consist of exons that are processed to produce mature RNAs and introns that are removed by RNA splicing. Alternative splicing is a regulated process by which exons can be either included or excluded in the mature RNA. Various RNAs (also called transcript variants) can be produced from a single gene through alternative splicing. Thus, alternative splicing enables a cell to express more RNA species than the number of genes it has, which increases the genome complexity^1^. In humans, for instance, approximately 95% of the multi-exonic genes undergoing alternative splicing have been revealed by high-throughput sequencing technology^2^. Exploring the functionality of alternative splicing is critical to our understanding of life mechanisms. Alternative splicing is associated with protein functions, such as diversification, molecular binding properties, catalytic and signaling abilities, and stability. Such related studies have been reviewed elsewhere^3,4^. Additionally, relationships between alternative splicing and disease^5^ or cancer^6,7^ have received increasing attention. Understanding the pathogenesis associated with alternative splicing can shed light on diagnosis and therapy. With the rapid development of high-throughput technology, it has become possible to study the transcriptome-wide function and mechanism of alternative splicing^8^.

The localization of RNA in a cell can determine whether the RNA is translated, preserved, modified, or degraded^9,10^. In other words, the subcellular location of an RNA is strongly related to its biological function^9^. For example, the asymmetric distribution of RNA in cells can influence the expression of genes^9^, formation and interaction of protein complexes^11^, biosynthesis of ribosomes^12^, and development of cells^13,14^, among other functions. Many techniques have been developed to investigate the subcellular localization of RNAs. RNA fluorescent in situ hybridization (RNA FISH) is a conventional method to track and visualize a specific RNA by hybridizing labeled probes to the target RNA molecule^15,16^. Improved FISH methods using multiplexing probes to label multiple RNA molecules have been presented, and have expanded the range of target RNA species^17,18^. With the development of microarray and high-throughput sequencing technologies, approaches for the transcriptome-wide investigation of RNA subcellular localizations have emerged^19^. Recently, a technology applying the ascorbate peroxidase APEX2 (APEX-seq) to detect proximity RNAs has been introduced^20,21^. APEX-seq is expected to obtain unbiased, highly sensitive, and high-resolved RNA subcellular localization *in vivo*. Simultaneously, many related databases have been developed^22,23^, which integrate RNA localization information generated by the above methods and provide valuable resources for further studies of RNA functions.

Alternative splicing can regulate RNA/protein subcellular localization^24–27^. However, to date, alternative splicing in only a limited number of genes has been examined. One approach to solve this problem involves the use of high-throughput sequencing. The goal of this research was to perform a comprehensive and transcriptome-wide study of the impact of alternative splicing on RNA subcellular localization. Therefore, we analyzed RNA-sequencing (RNA-seq) data obtained from subcellular (cytoplasmic and nuclear) fractions and investigated whether alternative splicing causes an imbalance in RNA expression between the cytoplasm and the nucleus. Briefly, we found that RNA splicing appeared to promote cytoplasmic localization. We also observed that transcripts with intron retentions preferred to localize in the nucleus. A further meta-analysis of retained introns indicated that short and structured intronic sequences were more likely to appear in nuclear transcripts. Additionally, motif analysis revealed that fourteen RBPs were enriched in the retained introns that are associated with nuclear localization. The above results were consistently observed across 13 cell lines, suggesting the reliability and robustness of the investigations. To our knowledge, this is the first transcriptome-wide study to analyze and evaluate the effect of alternative splicing on RNA subcellular localization across multiple cell lines. We believe that this research may provide valuable clues for further understanding of the biological function and mechanism of alternative splicing.

## 2 Materials and Methods

### 2.1 RNA-Seq Data and Bioinformatics

We used the RNA-seq data from the ENCODE^28^ subcellular (nuclear and cytoplasmic) fractions of 13 human cell lines (A549, GM12878, HeLa-S3, HepG2, HT1080, HUVEC, IMR-90, MCF-7, NHEK, SK-MEL-5, SK-N-DZ, SK-N-SH, and K562) to quantify the localization of the transcriptome. In brief, cell lysates were divided into the nuclear and cytoplasmic fractions by centrifugal separation, filtered for RNA with poly-A tails to include mature and processed transcript sequences (>200 bp in size), and then subjected to high-throughput RNA sequencing. Ribosomal RNA molecules (rRNA) were removed with RiboMinus™Kit. RNA-seq libraries of strand-specific paired-end reads were sequenced by Illumina HiSeq 2000. See Supplementary Data 1 for accessions of RNA-seq data. The human genome (GRCh38) and comprehensive gene annotation were obtained from GENCODE (v29)^29^. RNA-seq reads were mapped with STAR (2.7.1a)^30^ and quantified with RSEM (v1.3.1)^31^ using the default parameters. Finally, we utilized SUPPA2^32^ to calculate the differential usage (change of transcript usage, Δ*TU* ^33^) and the differential splicing (change of splicing inclusion, ΔΨ^34^) of transcripts.

### 2.2 Δ*TU* **and** ΔΨ

To investigate the inconsistency of subcellular localization of transcript variants, we calculated the Δ*TU* from the RNA-seq data to quantify this bias. For each transcript variant, Δ*TU* indicates the change in the proportion of expression level in a gene, which is represented as

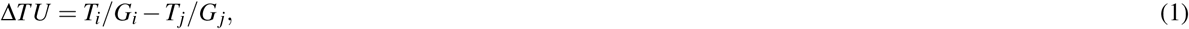

where *i* and *j* represent different subcellular fractions (i.e., cytoplasmic and nuclear, respectively). *T* and *G* are the expression abundance (transcripts per million, TPM) of the transcript and the corresponding gene, respectively. Expression abundances were estimated from RNA-seq data using the aforementioned RSEM. A high Δ*TU* value of a transcript indicates that it is enriched in the cytoplasm, and a low value indicates that it is enriched in the nucleus. We utilized SUPPA2 to compute Δ*TU* and assess the significance of differential transcript usage between the cytoplasm and the nucleus. For a gene, we selected a pair of transcript variants *T*_*c*_ and *T*_*n*_ that

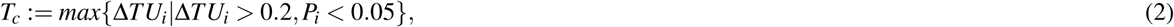

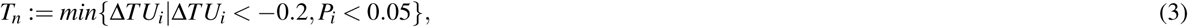

where *P*_*i*_ is the significance level of the *i*-th transcript variant. SUPPA2 calculates the significance level by comparing the empirical distribution of Δ*TU* between replicates and conditions (see^32^ for details). To study whether the different splicing patterns affect the subcellular localization of transcript, we used ΔΨ to quantify this effect. Seven types of splicing patterns— alternative 3′ splice-site (A3), alternative 5′ splice-site (A5), alternative first exon (AF), alternative last exon (AL), mutually exclusive exon (MX), retained intron (RI), and skipping exon (SE)—were considered. These splicing patterns were well-defined in previous studies^32, 35^. For each alternative splicing site (e.g., RI),

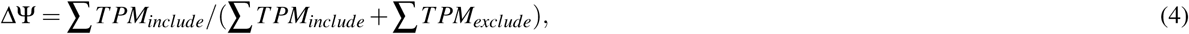

where *TPM*_*include*_ and *TPM*_*exclude*_ represent the expression values of transcripts included and excluded, respectively, from the splicing site.

### 2.3 Gene Ontology (GO) Enrichment Analysis

GO enrichment analysis was performed with g:Profiler (Benjamini–Hochberg false discovery rate (FDR) < 0.001)^36^. g:Profiler applies the hypergeometric test to detect statistically significantly enriched molecular function (MF), biological processes (BP), and cellular component (CC). In addition to GO enrichment analysis of switching genes in each cell line, we also defined the shared switching genes (detected as switching genes ≥ 5 cell lines in this study, *n* = 1219) and conducted GO enrichment analysis.

### 2.4 Nonsense-Mediated RNA Decay (NMD) Sensitivity

NMD sensitivity was defined by the fold-change of transcript expression before and after knockdown of *UPF1, SMG6*, and *SMG7* (main factors in NMD) in HeLa-S3 cells. The larger the value, the higher the degradation rate by NMD. We used the RNA-seq data (GSE86148)^37^ to calculate the NMD sensitivity. RNA-seq reads were mapped with STAR (2.7.1a)^30^ and quantified with RSEM (v1.3.1)^31^ using the default parameters and GENCODE (v29) comprehensive gene annotation. EBseq (v1.20.0)^38^ was used to calculate the fold-change of transcript expression.

### 2.5 RNA Structure Context Analysis

We used CapR^39^ with default parameters to predict secondary structures for RNA sequences. We extracted the stem probabilities of RNA sequence to represent the stability of the RNA secondary structure. The higher the stem probability, the more stable the secondary structure of RNA.

### 2.6 RBP-Binding Prediction and Repeat Sequence Analysis

We applied the stand-alone version of RBPmap^40^ to predict RBP binding sites. All RBPmap human/mouse stored motifs were used, and other parameters used default values. We used the repeat library (built on 31 January 2014) that mapped to human (hg38) from Repeatmasker^41^. Bedtools^42^ was used to obtain overlaps between these repeat features (TEs) and the target sequences (i.e., retained introns).

## 3 Results

### 3.1 Thousands of Transcript Switches between Cytoplasm and Nucleus Were Identified across Thirteen Human Cell Lines

To assess the influence of alternative splicing on RNA subcellular localization, we analyzed RNA-seq datasets that cover the nuclear and cytoplasmic fractions in 13 human cell lines (Supplementary Data 1). Using these datasets, we calculated transcript usage changes (Δ*TU*) to identify transcript switches for genes between the cytoplasm and the nucleus. For a transcript, the transcript usage (*TU*) means the percentage of its expression in all transcript variants, and the Δ*TU* assesses the extent of differential transcript usage between two conditions (i.e., nuclear and cytoplasmic fractions). Thus, a transcript switch involves two transcript variants in a gene, one of which is predominantly expressed in the cytoplasm (Δ*TU >* 0, termed “cytoplasmic transcripts”), while the other one is mainly expressed in the nucleus (Δ*TU <* 0, termed “nuclear transcripts”). For instance, the *C16orf91* gene contains two transcript variants (Figure 1A, upper). In HeLa-S3 cells, we observed that the *C16orf91-201* and *C16orf91-202* (also termed a transcript switch in this study) were prone to substantial expression in the cytoplasm and the nucleus, respectively (see Figure 1A, lower and Supplementary Data 2). In total, 2073 transcript switches were detected in HeLa-S3 cells (Figure 1B). Specially, using a *p*-value of <0.05, we further screened for the switches that were not significantly altered. For example, in HT1080, SK-MEL-5, and SK-N-DZ, we appropriately discarded transcripts that were more likely to exhibit changes in transcript usage because of low expression levels (Supplementary Data 3).

**Figure 1.**
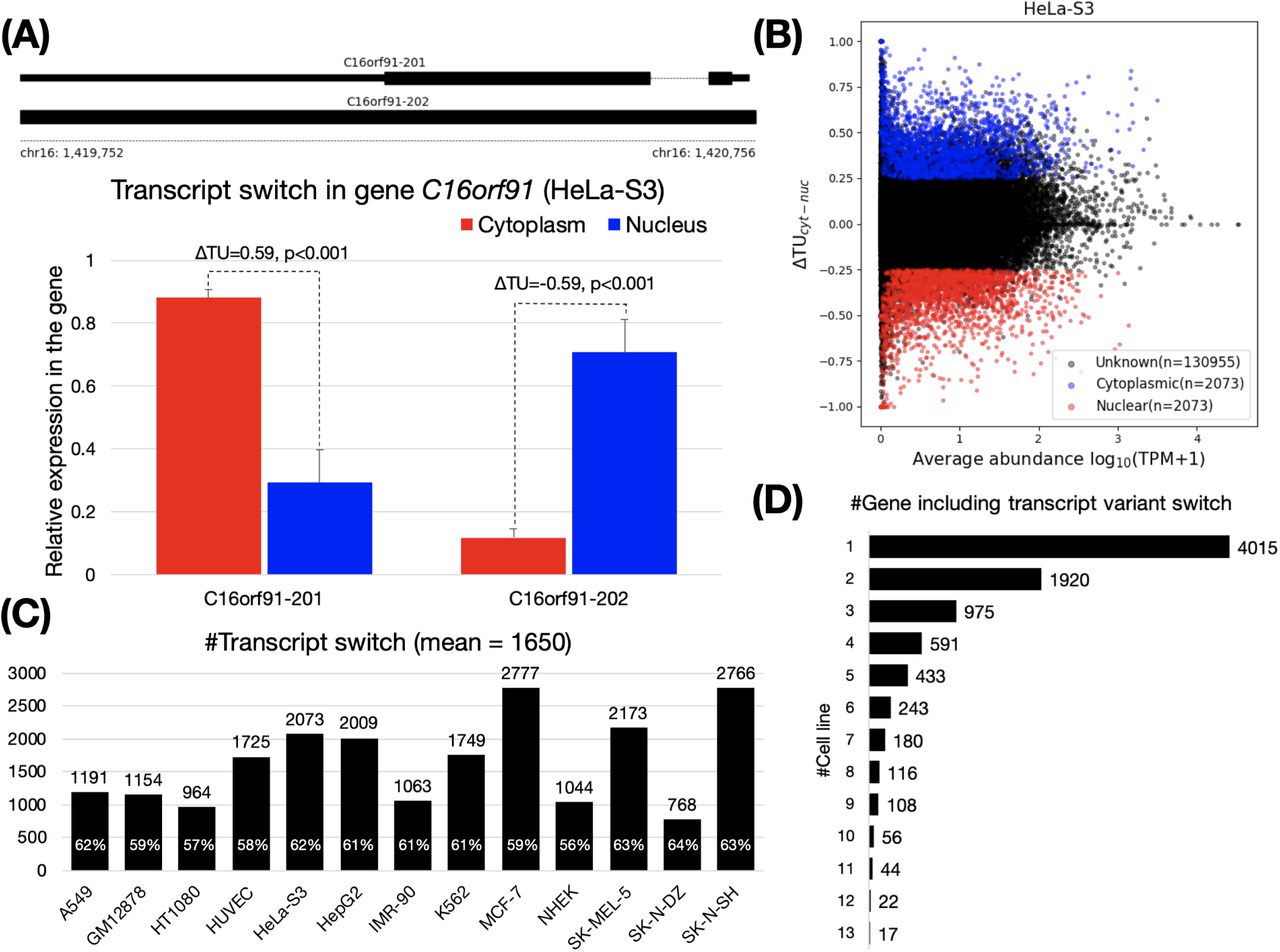
Transcript switches between cytoplasm and nucleus. (**A**) The upper panel shows two transcript variants of the gene *C16orf91*. The lower panel is an example for the identification of a transcript variant switch in the gene *C16orf91* using Δ*TU*. C16orf91-201 and C16orf91-202 were mainly localized in the cytoplasm and the nucleus, respectively, in HeLa-S3 cells (see Supplementary Data 2 for read coverage on *C16orf91* gene). P values are calculated with SUPPA2^32^ (see Materials and Methods). (**B**) Transcriptome-wide identification of transcript variant switches in HeLa-S3 cells. We applied |Δ*TU*| *>* 0.25 and *p <* 0.05 to filter out cytoplasmic (|Δ*TU*| *>* 0, blue) and nuclear (|Δ*TU*| *<* 0, red) transcript variants. Unknown (black) are transcripts filtered as no significant. (**C**) Number of transcript variant switches across the 13 cell lines. The percentage on the histogram indicates the content of protein-coding transcript. (**D**) Number of genes containing transcript variant switches shared across the 13 cell lines. For example, 4015 genes were identified as being cell specific.

On average, 1650 pairs of transcript switches were identified across the 13 cell lines (Figure 1C). The smallest (768 pairs) and the largest (2766 pairs) were both observed in brain tissue, SK-N-DZ and SK-N-SH cells, respectively. Next, we assessed if there is a group of genes shared among cell lines with similar biological functions through the inclusion of transcript switches. We observed a total of 8720 genes containing transcript switches from these 13 cell lines (hereafter referred to as “switching genes”). More than 93% (8177 of 8720) of these switching genes were found in fewer than six cell lines, indicating that most switching genes were cell-specific (Figure 1D and Supplementary Data 4, some may also caused by the cell specificity of the gene expression). GO enrichment analysis of the remaining genes revealed that these shared switching genes (detected as switching genes ≥ 5 cell lines, *n* = 1219) are associated with receptor/transducer activities (adjusted *p* < 1 × 10^−10^, GO:0004888, GO:0060089, GO:0038023, and GO:0004930), multicellular organismal process and G protein-coupled receptor signaling pathway (adjusted *p* < 1 × 10^−10^, GO:0032501 and GO:0007186), plasma membrane and cell periphery (adjusted *p* < 1 × 10^−10^, GO:0005886 and GO:0071944). Although the GO enrichment analysis results of switching genes in each cell line are somewhat different, they are also roughly similar. (see Supplementary Data 5).

To validate the transcripts defined by Δ*TU*, we first compared the Δ*TU* between protein-coding and non-coding transcripts over cell lines. Consistently, many cytoplasmic transcripts were observed to be protein-coding transcripts with positive Δ*TU* values, while non-coding transcripts preferred to locate in the nucleus and displayed more negative Δ*TU* values (all *p* < 5.4 × 10^−24^, Supplementary Data 6). This result agreed with our understanding that the protein-coding transcript needs to be transported into the cytoplasm to produce proteins, while a large number of non-coding transcripts have been reported to localize in the nucleus to participate in transcriptional and post-transcriptional gene expression and chromatin organization, among other functions^43, 44^.

### 3.2 A Transcript Switch Is Associated with RNA Splicing rather than RNA Degradation

We first investigated the post-transcriptional influence of the NMD pathway on the transcript switch. Initially, the two transcripts in a transcript switch are equally distributed in the cytoplasm and nucleus. We consider one possible reason for the transcript switch is that the rate of degradation of the two transcripts in the cytoplasm and nucleus is different. To test this hypothesis, we calculated the sensitivity of transcripts to NMD in HeLa-S3 cells based on changes in RNA-seq data^37^ before and after the knockout of *UPF1, SMG6*, and *SMG7* (core factors of NMD). However, after comparing the sensitivity of NMD to cytoplasmic and nuclear transcripts, we did not find significant differences (Figure 2A). This observation led us to conclude that NMD may not significantly affect transcript switches. The cause of the transcript switch should be mainly due to intrinsic (sequence features) and transcriptional (e.g., splicing) effects.

**Figure 2.**
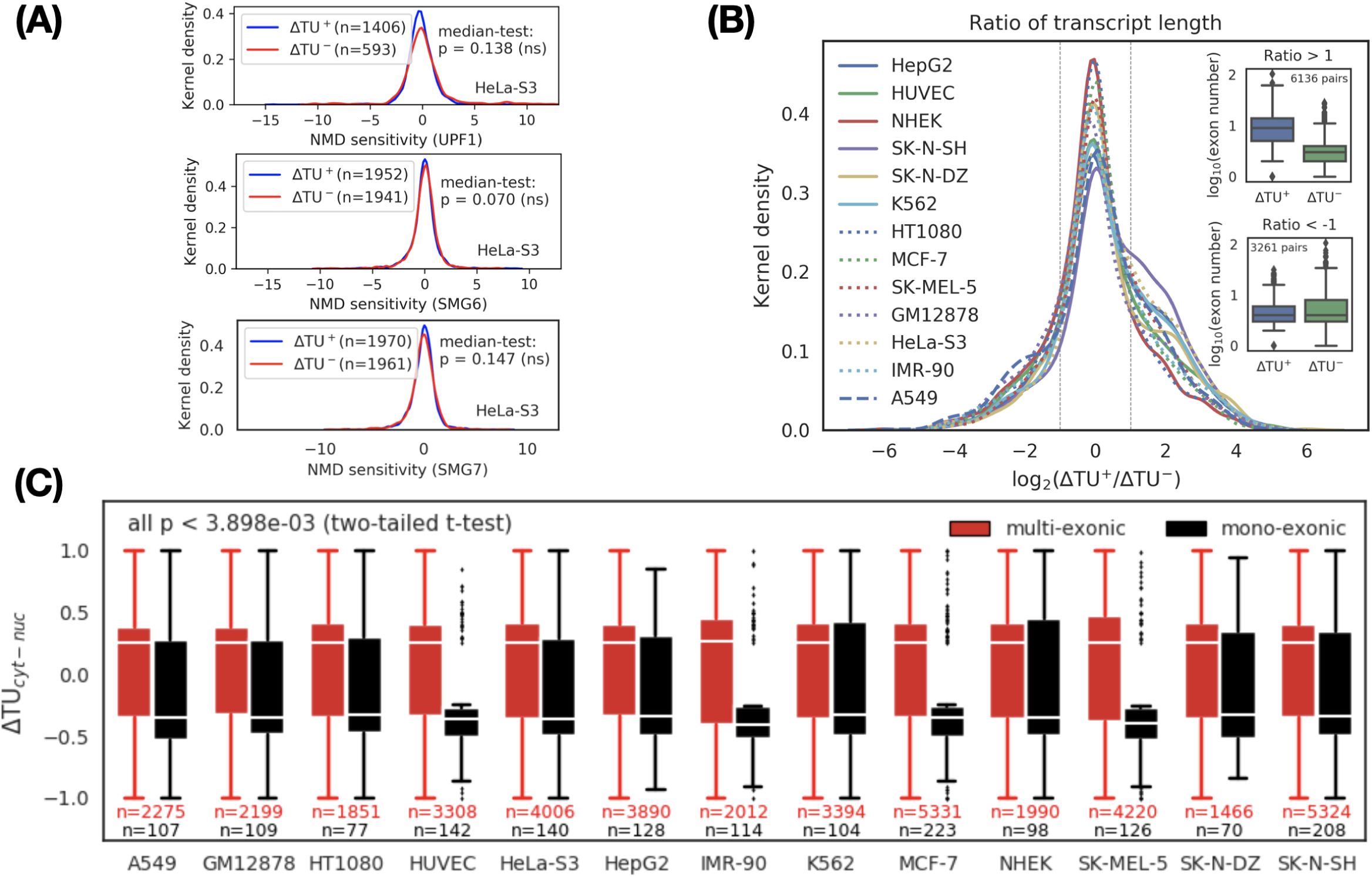
Characterization of cytoplasmic and nuclear transcripts. (**A**) NMD does not explain transcript variant switches between the cytoplasm and the nucleus. NMD sensitivity was defined by the fold-change of transcript expression before and after knockdown of NMD factor *UPF1* (upper), *SMG6* (middle), and *SMG7* (lower) in HeLa-S3 cells. P values: Mood’s median test. ns: not significant. (**B**) Comparison of transcript length between cytoplasmic (Δ*TU* ^+^) and nuclear (Δ*TU*^−^) transcripts. For each pair of transcript switches, the ratio of transcript length between the cytoplasm and the nucleus is defined as a metric. *Ratio >* 1 indicates a group of transcripts where the length of the cytoplasmic transcripts is greater than that of the nuclear transcripts. In the group, splicing frequency (or exon number) is over-expressed in the cytoplasmic transcripts (inset, upper, *p* < 2.2 × 10^−308^ Mood’s median test), suggesting splicing can promote cytoplasmic localization. No significant difference in splicing frequency was observed in the group if the transcripts *Ratio <* − 1 (inset, bottom, *p* < 4.8 × 10^−12^ Mood’s median test). (**C**) Comparison of Δ*TU* between mono-exonic (black) and multi-exonic (red) transcripts across 13 cell lines. Δ*TU* shows a significant positive correlation with splicing, indicating that splicing appears to be a dominant factor for RNA export from the nucleus. Based on two-tailed t-test, we calculated the significant level of difference between Δ*TU* in mono-exonic and multi-exonic transcripts in each cell line. The largest p value is 3.898 × 10^−3^.

Considering that the longer the transcript is, the higher the splicing frequency (or exon number) should be^45^, we next determined the association between the length of the transcript and its subcellular localization. We evaluated the relationship between length and subcellular localization by calculating the length ratio (logarithmic scale) between the cytoplasmic transcript and the nuclear transcript in each transcript switch. We divided the transcript switches into three categories based on the length ratio: positive (*ratio >* 1), negative (*ratio <* − 1), and neutral (other). A positive category indicates that the longer the transcript, the more enriched in the cytoplasm, and vice versa. A neutral category means that there is weak or no correlation (|*ratio*| ≤ 1) between transcript length and its subcellular localization. We observed that the number of transcript switches in the positive category was higher than that in the negative category (*p* < 1 × 10^−6^, Binomial test), which implied that the longer the transcript is, the more likely it is to be transported into the cytoplasm (Figure 2B). To verify whether the splicing frequency of the transcript is positively correlated with its cytoplasmic location, we further compared the distribution of exon numbers (i.e., splicing frequency) in the cytoplasmic transcripts and nuclear transcripts for the positive and negative categories, respectively (Figure 2B, inset). We found that the exon number of the cytoplasmic transcripts in the positive category was higher than the nuclear transcripts (see Figure 2B, inset upper, *p* < 2.2 × 10^−308^ Mood’s median test). Based on the above observations, we speculated that there are significant differences in subcellular localization between transcripts with or without splicing events. To confirm this hypothesis, we divided the transcript switches into the mono-exonic (unspliced) and multi-exonic (spliced) groups and then compared the distribution of Δ*TU* values between them. As expected, the Δ*TU* value of multi-exonic transcripts was positive (indicating cytoplasmic localization) and was significantly and consistently higher than the negative Δ*TU* value of mono-exonic transcripts (representing nuclear localization) in all cell lines (Figure 2C and Supplementary Data 7).

### 3.3 Enrichment and Characterization of Retained Introns in the Nuclear Transcripts

Next, we asked whether there was a specific kind of splicing pattern associated with subcellular localization. Seven different splicing patterns (A3, A5, AF, AL, MX, RI, and SE) were considered^46^. Comparing to an original transcript, each type of splicing pattern inserts, deletes, or replaces a partial sequence that may include sequence elements, which have important or decisive effects on subcellular localization (e.g., protein-binding sites, RNA structures, etc.). We used the ΔΨ, which ranges from − 1 to 1, to measure the preference for a type of splicing pattern between the nucleus and the cytoplasm (see “Materials and Methods” for details). The smaller the ΔΨ, the more the corresponding splicing pattern prefers the nucleus. Interestingly, in HeLa-S3 cells, we observed that retained introns were biased toward the nucleus, while other splicing patterns had no preference (Figure 3A). Consistent results were also observed in the other twelve cell lines (Supplementary Datas 8 and 9).

**Figure 3.**
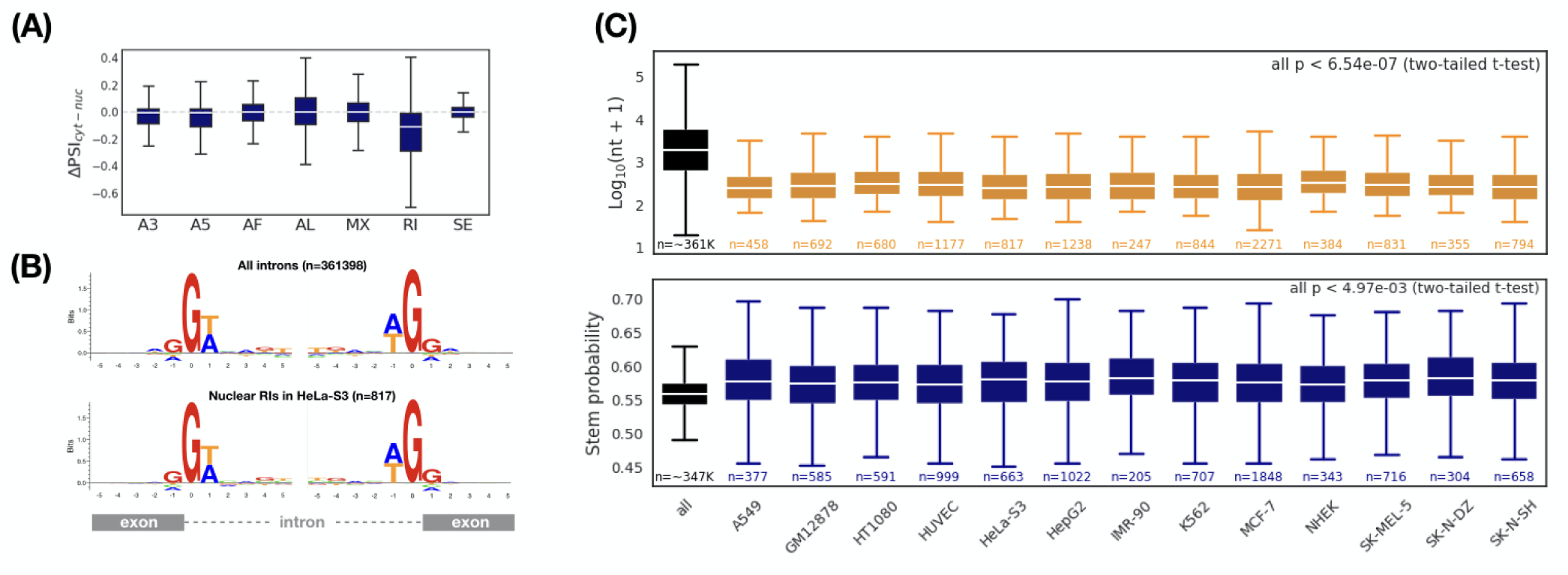
Comparison of alternative splicing patterns between cytoplasmic and nuclear transcripts. (**A**) Retained introns are enriched in the nuclear transcripts (HeLa-S3). ΔΨ indicates the inclusion level for a specific alternative splicing pattern. A3: alternative 3 splice-site; A5: alternative 5 splice-site; AF: alternative first exon; AL: alternative last exon; MX: mutually exclusive exon; RI: retained intron; SE: skipping exon. Comparison of (**B**) splice sites, (**C**) Length (top) and RNA secondary structure (bottom) between all introns and nuclear RIs (ΔΨ *<* 0 and *p <* 0.05). Sequence logos show intron-exon (the left) and exon-intron (the right) splice boundaries. *p* values: two-tailed *t*-test between all introns and nuclear RIs. The largest *p* value is 6.54 × 10^−7^ and 4.97 × 10^−3^ for length and RNA secondary structure, respectively.

To further characterize the retained introns associated with nuclear localization, we first selected a set of retained introns enriched in the nucleus (ΔΨ *<* 0 and *p <* 0.05, termed nuclear RIs). Compared with all retained introns, we should be able to observe some different features of the nuclear RIs, which are likely to be the determining factors for nuclear localization. We first investigated whether the nuclear RIs have a specific splicing signal. By applying this splicing signal, it is possible to regulate the intron retention specifically, thereby monitoring the subcellular localization of the transcript. Unfortunately, we found no differences in the splicing signal between the nuclear RIs and all RIs (Figure 3B). We then considered whether the structure of the nuclear RIs (including the length and RNA secondary structure) have a specific signature. Surprisingly, the length of the nuclear RIs was significantly shorter than the overall length level (Figure 3C, upper (all *p* < 6.64 × 10^−7^) and Supplementary Data 10). We also found that the average stem probability of the nuclear RIs was higher than that of all RIs (Figure 3C, bottom and Supplementary Data 11). Thus, we concluded that nuclear RIs maintain a more compact and stronger structure when compared with the overall RIs.

### 3.4 The Preferences of RNA-Binding Proteins on Retained Introns Suggest Their Role in Nuclear Localization

We sought to further investigate whether the nuclear RIs mentioned above are associated with RBPs. We calculated the frequency of each dinucleotide in the nuclear RIs across the 13 cell lines and normalized it with the background frequency of all intronic sequences in the genome. The reason we examined the dinucleotide composition of nuclear nucleotides was due to the reported specific contact between amino acids and dinucleotides^47^. The normalized dinucleotide frequency represents the extent to which the corresponding dinucleotide is preferred in the nuclear RIs. Overexpression of the GC-rich sequence occurred in the nuclear RIs (Figure 4A). This observation also provided an intuitive hypothesis of whether RBPs that preferentially bound to RNA containing GC-rich sequences directly or indirectly affect subcellular localization.

**Figure 4.**
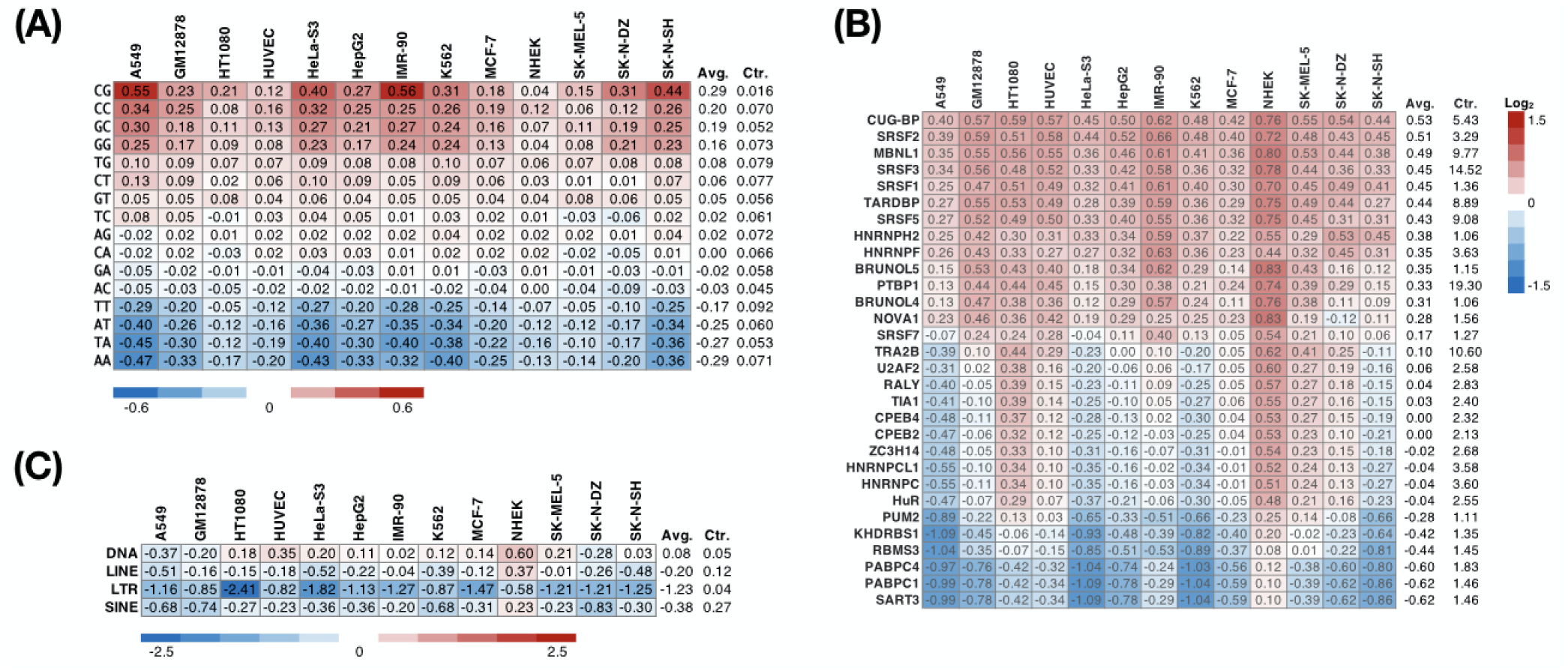
Preference of dinucleotide, RBPs, and TEs on nuclear RIs. The heatmap represents the relative density of a specific (**A**) dinucleotide, (**B**) RBP-binding site, or (**C**) TE (row) on nuclear RIs across multiple cell lines (columns) compared with that of all introns (Ctr.).

To further explore which RBPs preferentially interact with nuclear RIs, we predicted the binding preference between an RBP and an intronic sequence using RBPmap^40^, a motif-based approach. We found that across the 13 cell lines, 14 of 30 RBPs tested consistently (more than 80%) preferred attaching to the nuclear RIs (Figure 4B). These include serine/arginine-rich proteins (SRSF1, SRSF2, SRSF3, SRSF5, SRSF7), heterogeneous nuclear ribonucleoproteins (HNRNPH2, HNRNPF, PTBP1), CUG-binding proteins (CUG-BP, MBNL1, BRUNOL4, BRUNOL5), and others (TARDBP and NOVA1). Conversely, binding motifs of five RBPs (SART3, PABPC1, PABPC4, RBMS3, KHDRBS1) were consistently under-represented in the nuclear RIs. Considering a previous report showing that the repeat sequence drives the nuclear localization of lncRNA^48^, we subsequently analyzed the enrichment of the repeat sequence in the nuclear RIs. In NHEK cells, we observed SINE (short interspersed nuclear element; consistent with the previous study^48^), LINE (long interspersed nuclear element), and DNA transposon enrichment in nuclear RIs (Figure 4C). However, such phenomena have not been seen in other cell lines, suggesting that retained introns may not be ubiquitous in guiding nuclear localization through repeat sequences. Additionally, we observed that an LTR (long terminal repeat) was relatively rare in the nuclear RIs in all cell lines, implying that an LTR is involved in the regulation of RNA subcellular localization.

## 4 Discussion

We emphasize that Δ*TU* measures the change in the proportion of gene expression for a transcript between the nucleus and the cytoplasm. An increased Δ*TU* does not imply that the transcript abundance in the cytoplasm is higher than that in the nucleus. Instead, a significantly increased Δ*TU* indicates that the transcript is stored dominantly for the corresponding gene in the cytoplasm, but is expressed as a minor isoform in the nucleus. Therefore, Δ*TU* measures the dynamic changes of a representative (dominantly expressed) transcript of a gene between the nucleus and the cytoplasm. By using Δ*TU*, we defined a transcript switch, which indicates that a single gene has two different representative transcripts in the nucleus and the cytoplasm, respectively. A gene containing a transcript switch is termed as a switching gene in this study. Due to the limitation of available experimental data, we calculated Δ*TU* and performed alternative splicing analysis based on short-read RNA-seq. However, long-read RNA-seq and direct RNA-seq can provide more clear and convincing splicing events^49–52^.

We hypothesize that a switching gene can function separately in the nucleus and cytoplasm (hereafter “bifunctional gene”), respectively. Previous studies have discovered several bifunctional mRNAs, which generate coding and non-coding isoforms through alternative splicing. The coding isoform translates proteins in the cytoplasm while the non-coding isoform resides in the nucleus to function as a scaffolding agent^53–57^, post-transcriptional component^58^, or coactivator^59–61^. Furthermore, a controlling feedback loop between two such isoforms can be considered. This hypothesis is consistent with a phenomenon called genetic compensation response (GCR), which has been proposed and validated in recent years^62–65^. In GCR, RNA sequence fragments obtained after mRNA degradation are imported to the nucleus to target genes based on sequence similarity and regulate their expression level. We argue that GCR-like feedback may be widely present in the switching genes. The reason is that parts of exonic sequences are usually shared (implying high sequence similarities) among transcript variants in the switching genes. Indeed, we have observed 768 ∼ 2777 switching genes (Figure 1C) in the 13 cell lines, suggesting that bifunctional genes or GCR are more widespread in the human genome than expected. A total of 8720 switching genes (Figure 1D) were identified in this study. We believe that this analysis promises to provide a valuable resource for studying bifunctional or GCR-associated genes. Additionally, we observed that most of the switching genes were cell-specific (Figure 1D), suggesting that such genes may be related to genetic adaption to the environmental changes (stress, development, disease, etc.).

Based on the observation that multi-exonic transcripts are preferentially exported to the cytoplasm, we argue that splicing supports cytoplasmic localization. One possible reason is that the exon-exon junction complex (EJC) may promote RNA export. The EJC containing or interacting with the export factor binds to the transcript during splicing, while the export factor promotes cytoplasmic localization. Previous studies have reported that the EJC component in *Xenopus* provides binding sites^66^ for export factors (REF and/or TAP/p15). The coupling between a conserved RNA export machinery and RNA splicing has also been reviewed^67^.

Additionally, we observed that transcript variants containing retained introns were more likely to reside in the nucleus, which is consistent with the previous studies^68, 69^. By further analyzing these retained introns, we found that transcripts, including short but structurally stable retained introns, were preferentially localized in the nucleus. It is worth noting that the above observations are still robust after we distinguish between protein coding and noncoding genes (Supplementary Datas 7, 10 and 11). We speculate that such introns function as platforms for attaching to localization shuttles (e.g., RBPs). Short introns appear to be favored by natural selection because of the low metabolic cost of transcription and other molecular processing activities^67^. The RNA structure context was discussed as being able to affect protein binding^70^. Motif analysis of the retained introns of nucleus-localized transcripts detetced 14 enriched RBPs, in which MBNL1 has been reported to bind RNA in the CUG repeats^71^ and contain two nuclear signals^72^. Previous studies have reported that the CUG repeat expansion caused nuclear aggregates and RNA toxicity, which in turn induces neurological diseases^73, 74^. Taken together, the data reveal a post-transcriptional regulation mechanism in which alternative splicing regulates RNA subcellular localization by including or excluding those introns containing nuclear transporter binding sites.

## 5 Conclusions

This study explored whether alternative splicing can regulate RNA subcellular localization. The RNA-seq data derived from the nuclear and cytoplasmic fractions of 13 cell lines were utilized to quantify transcript abundance. We applied Δ*TU* to define a pair of nuclear or cytoplasmic transcripts expressed from a single gene locus. By comparing nuclear and cytoplasmic transcripts, we consistently observed that splicing appears to promote RNA export from the nucleus^75^. Furthermore, we used ΔΨ to analyze the effect of splicing patterns on RNA localization and found that intron retention was positively correlated with nuclear localization. Sequence analysis of the retained introns revealed that short and structurally stable introns were favored for nuclear transcripts. We argue that such intronic sequences provide a hotspot for interacting with nuclear proteins. Subsequently, fourteen RBPs are consistently identified to prone to associate with retained introns across 13 cell lines. We argue that cells can regulate the subcellular localization and biological functions of RNAs through alternative splicing of introns, which contain localization elements (RNA structure context or sequence motifs). These findings can help us better understand the regulatory mechanisms of RNA localization, and thus, for example, predict RNA localization more precisely from sequence characteristics^76^.

Although we used an EM-based method^31^ to predict the transcript abundance between overlapping transcripts, this transcript quantification is still very challenging. One possible solution is to obtain more accurate transcript structures and their expression level through long reads (such as Nanopore sequencing^77, 78^). Studying the effects of alternative splicing on RNA localization in healthy tissues or other species would be another challenging work. In summary, this research provides important clues for studying the mechanism of alternative splicing on gene expression regulation. We also believe that the switching genes defined in this study (Supplementary Data 4) will provide a valuable resource for studying gene functions.

## Acknowledgements

C.Z. and M.H. are grateful to Martin Frith, Yutaka Saito, and Masahiro Onoguchi for valuable discussions. Computations were partially performed on the NIG supercomputer at ROIS National Institute of Genetics. This work was supported by the Ministry of Education, Culture, Sports, Science and Technology (KAKENHI) [grant numbers JP17K20032, JP16H05879, JP16H01318, and JP16H02484 to M.H.].

## Author contributions

M.H. and C.Z. conceived this study and contributed to the analysis and interpretation of the data. C.Z. performed all the experiments and wrote the draft, M.H. revised it critically. All authors read and approved the final manuscript.

## Additional information

- Supplementary Data 1 — RNA-seq data used in this study.
- Supplementary Data 2 — Read coverage of gene *C16orf91* in HeLa-S3.
- Supplementary Data 3 — Transcriptome-wide identification of transcript variant switches in 12 human cell lines.
- Supplementary Data 4 — List of switching genes identified across 13 cell lines in this study. Gene Ontology enrichment analysis for switch genes.
- Supplementary Data 5 — Gene Ontology enrichment analysis for switch genes.
- Supplementary Data 6 — Comparison of Δ*TU* between protein-coding transcripts and non-coding transcripts.
- Supplementary Data 7 — Comparison of Δ*TU* between mono-exonic and multi-exonic transcripts across 13 cell lines.
- Supplementary Data 8 — Comparison of alternative splicing patterns between cytoplasmic and nuclear transcripts (all genes).
- Supplementary Data 9 — Comparison of alternative splicing patterns between cytoplasmic and nuclear transcripts (protein-coding and noncoding genes).
- Supplementary Data 10 — Comparison of length between all introns and nuclear RIs.
- Supplementary Data 11 — Comparison of RNA secondary structure (stem probability) between all introns and nuclear RIs.

**Supplementary Data 1.**
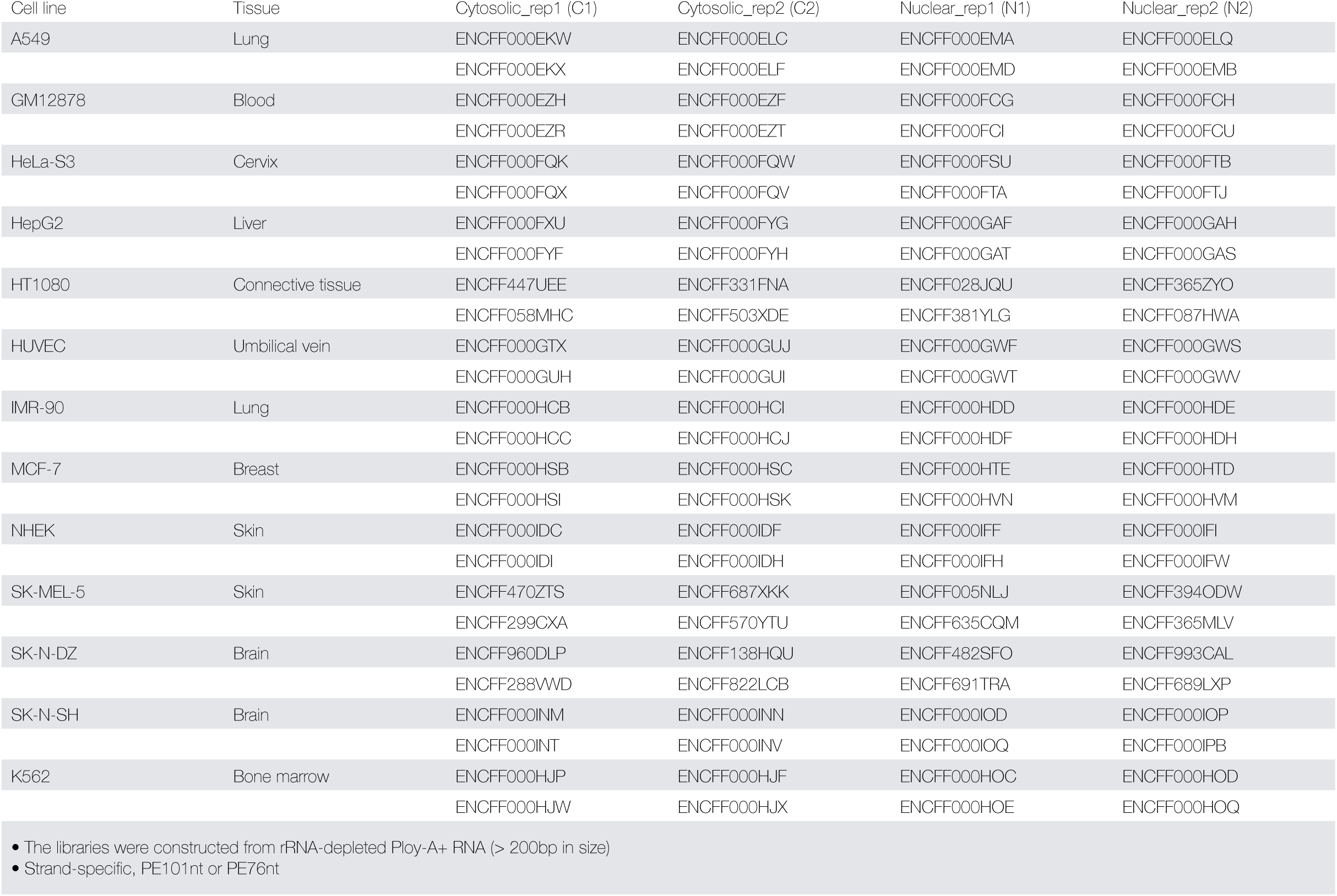
RNA-seq data used in this study

**Supplementary Data 2.**
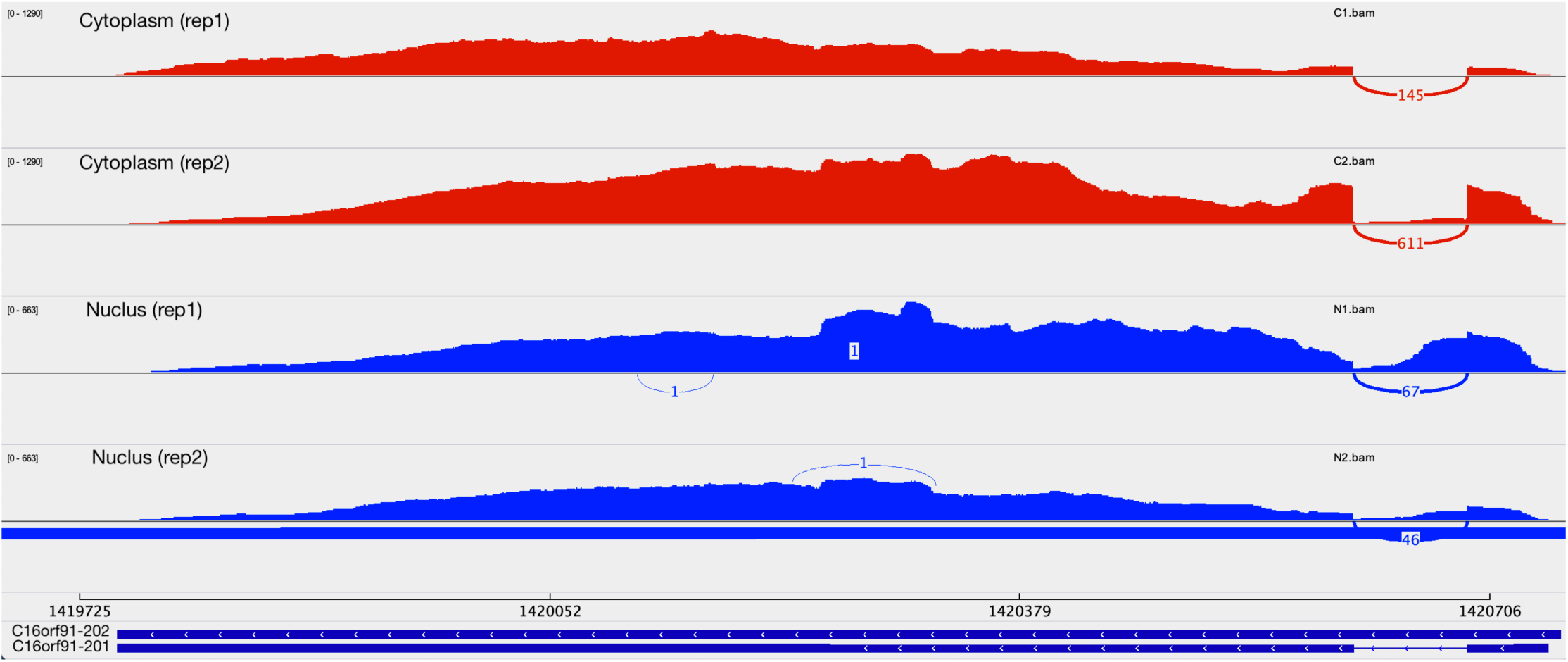
Read coverage of gene *C16orf91* in HeLa-S3. IGV (Robinson et al 2011) was used to visualize the read coverage.

---

**Supplementary Data 3.**
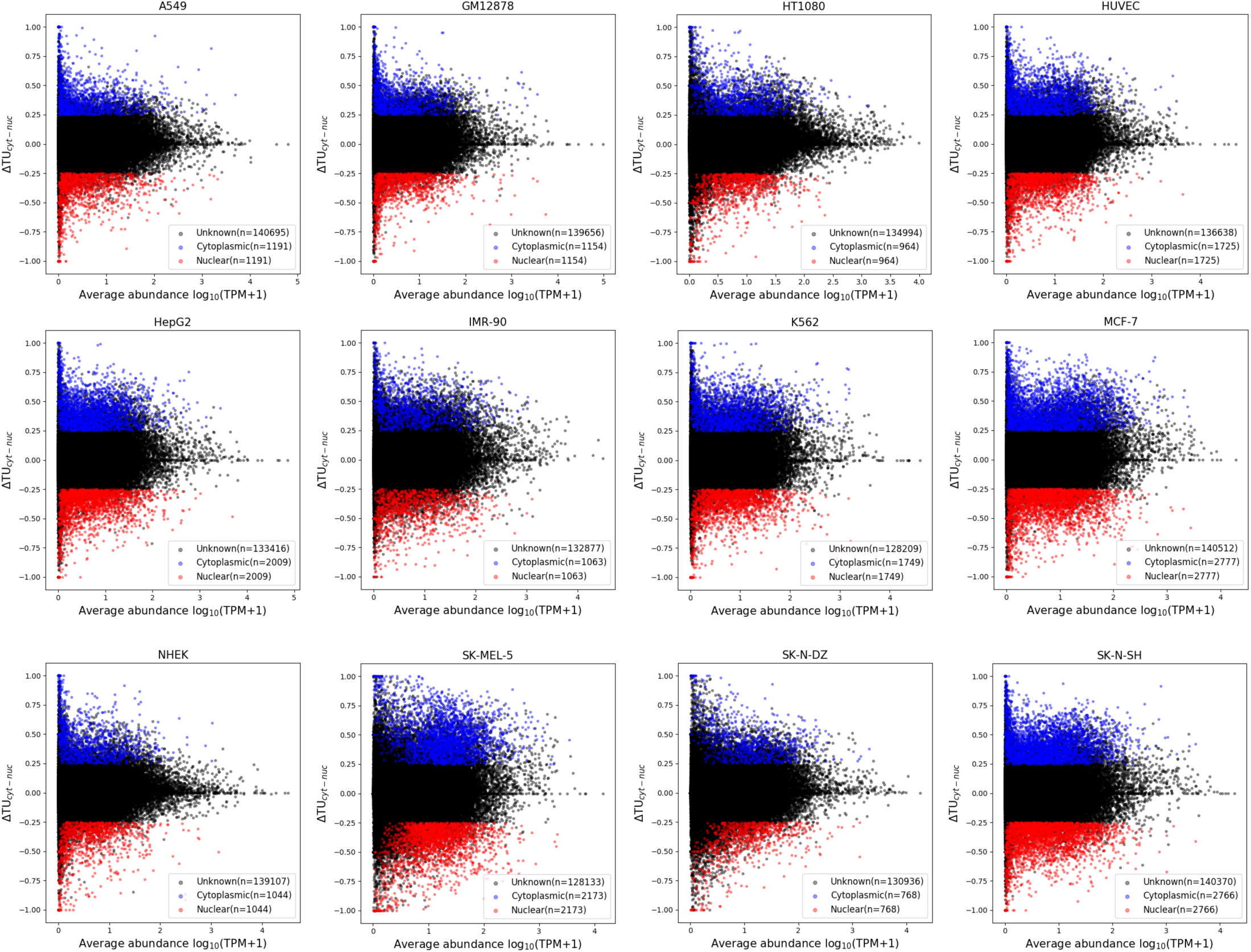
Transcriptome-wide identification of transcript variant switches in 12 human cell lines. We applied |ΔTU|>0.25 and p<0.05 to filter out cytoplasmic(|ΔTU|>0, blue) and nuclear (|ΔTU|<0, red) transcript variants. Unknown (black) are transcripts filtered as no significant.

**Supplementary Data 4. List of switching genes identified across 13 cell lines in this study.** Online resource: https://doi.org/10.6084/m9.figshare.11999658

**Supplementary Data 5.**
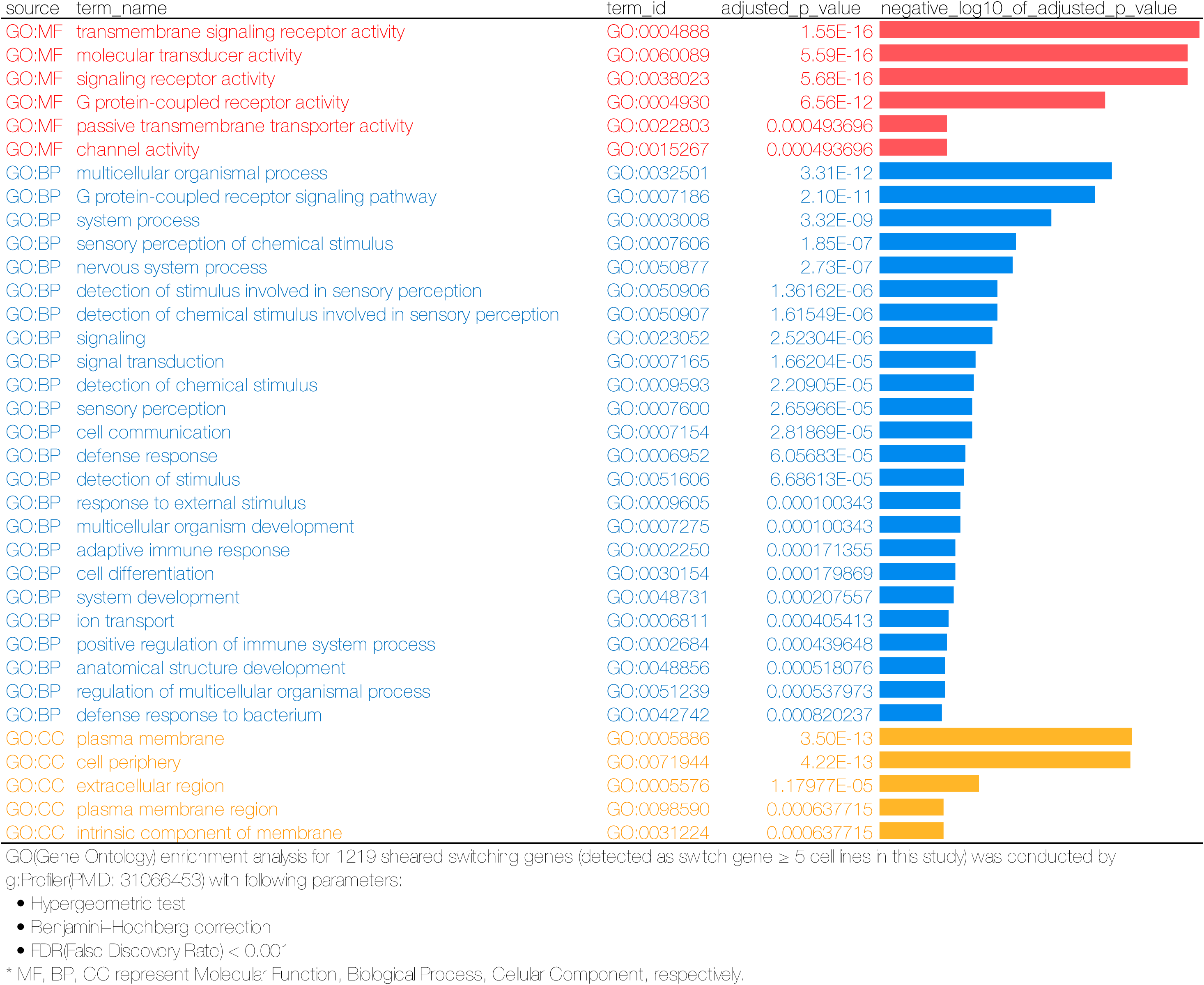
Gene Ontology enrichment analysis for switch genes (sheared, continued)

**Supplementary Data 5.**
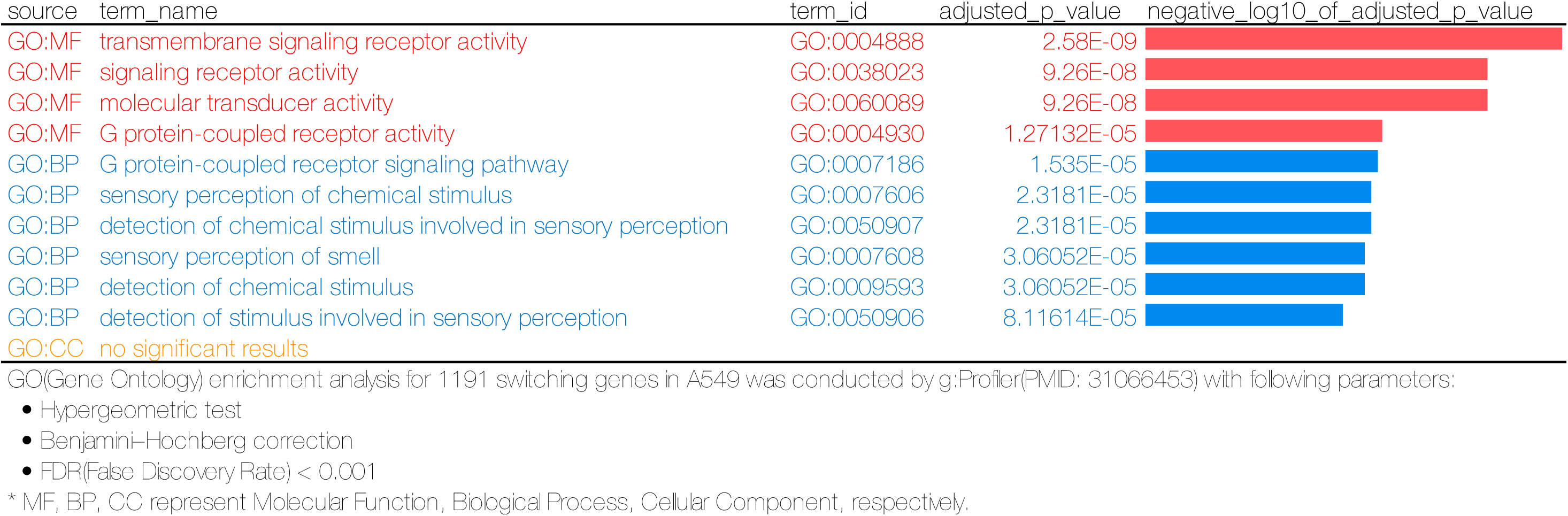
Gene Ontology enrichment analysis for switch genes (A549, continued)

**Supplementary Data 5.**
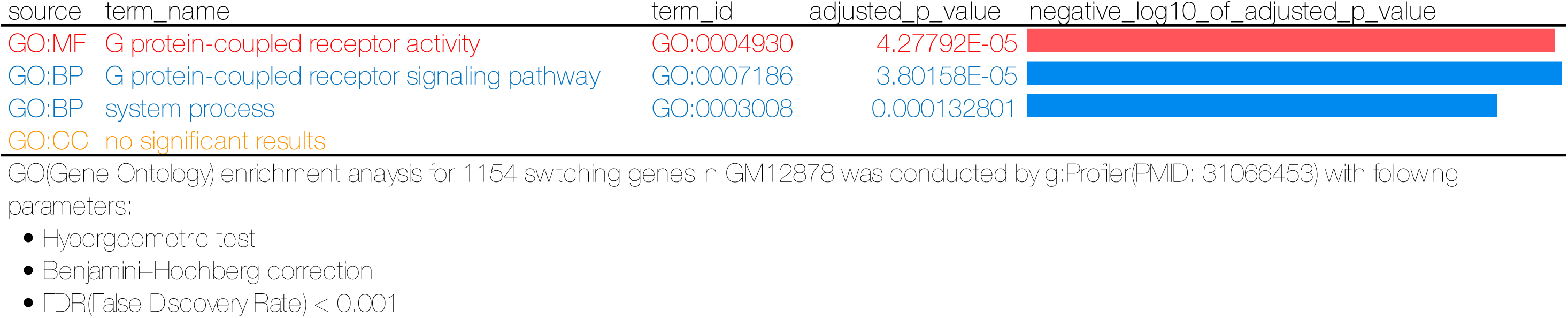
Gene Ontology enrichment analysis for switch genes (GM12878, continued)

**Supplementary Data 5.**
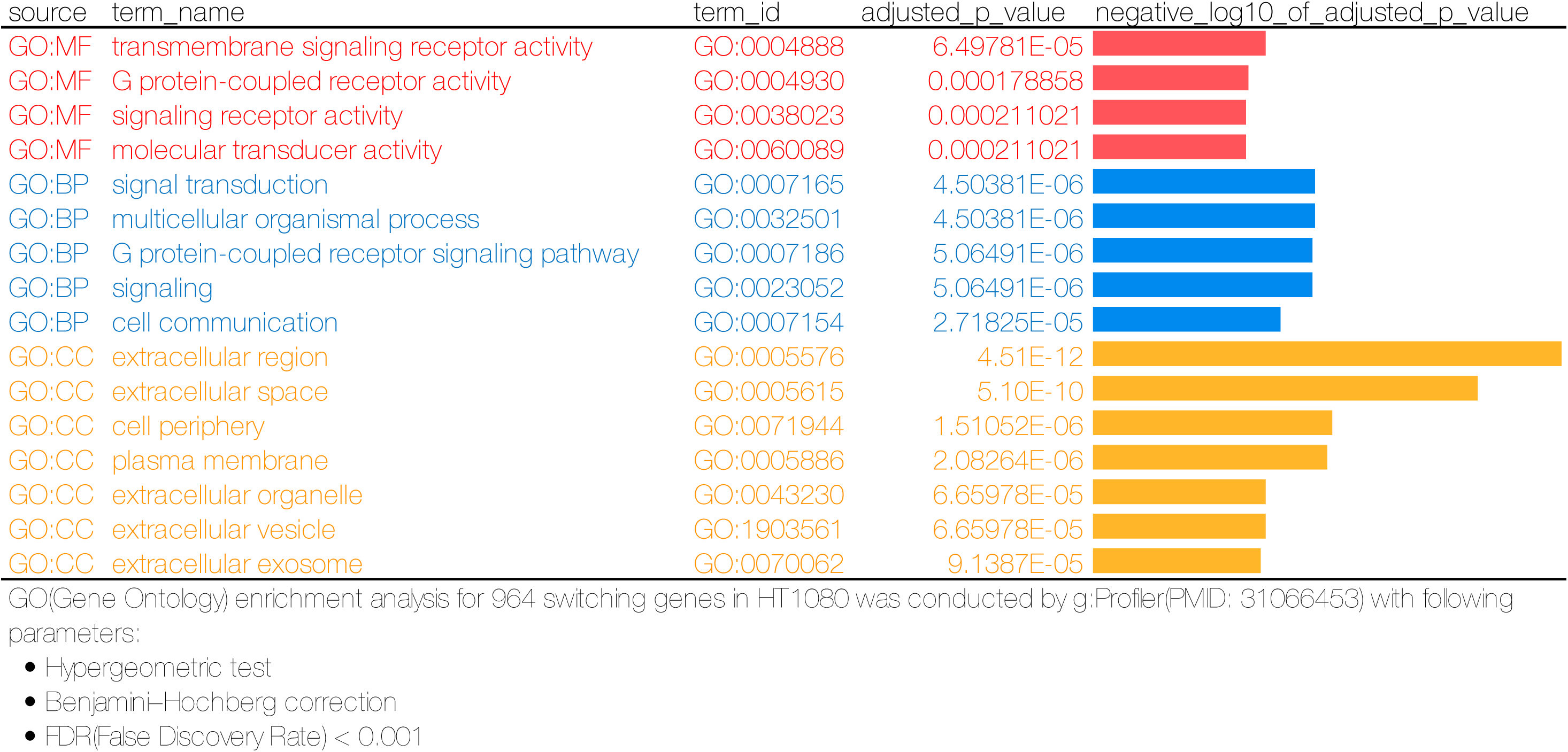
Gene Ontology enrichment analysis for switch genes (HT1080, continued)

**Supplementary Data 5.**
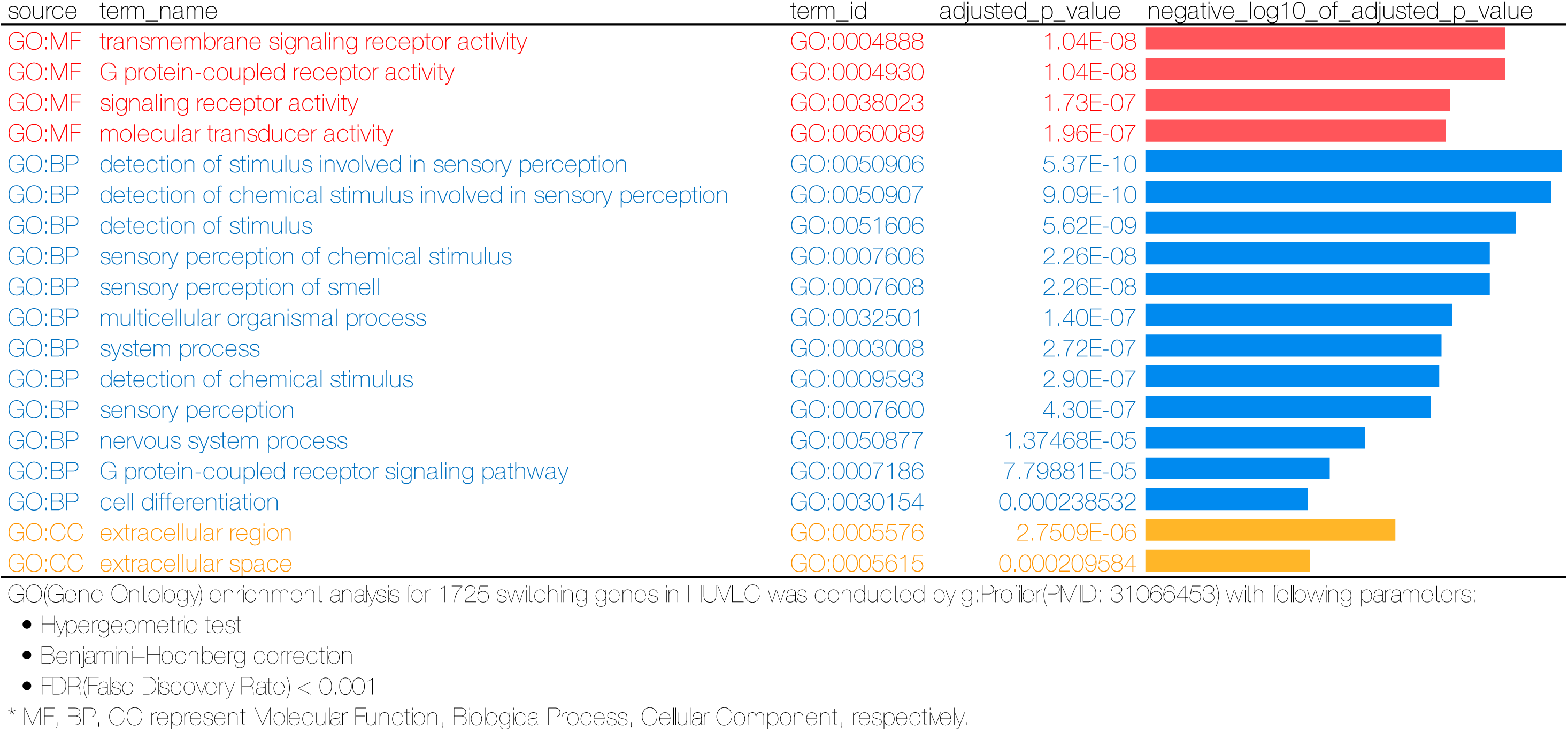
Gene Ontology enrichment analysis for switch genes (HUVEC, continued)

**Supplementary Data 5.**
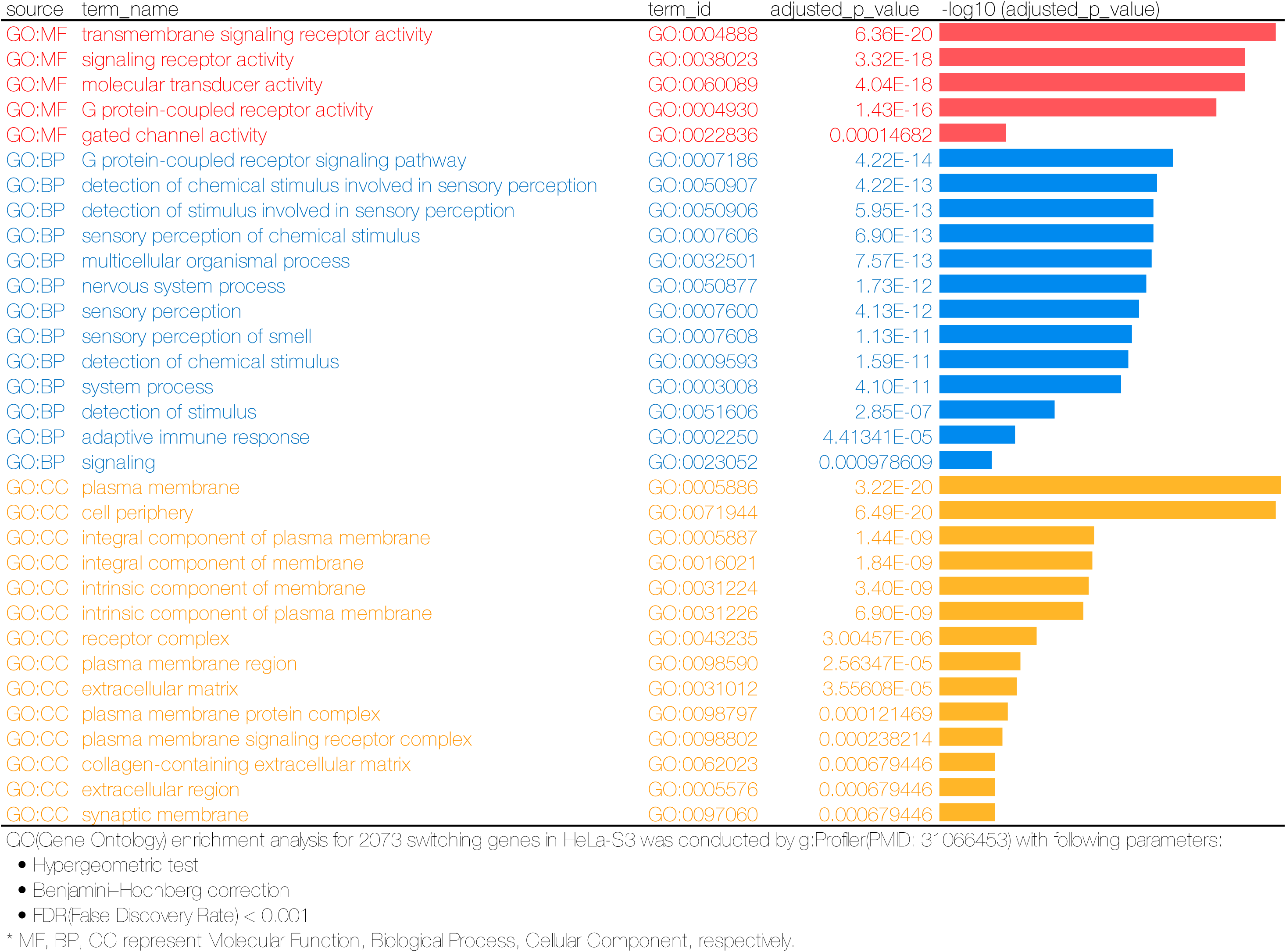
Gene Ontology enrichment analysis for switch genes (HeLa-S3, continued)

**Supplementary Data 5.**
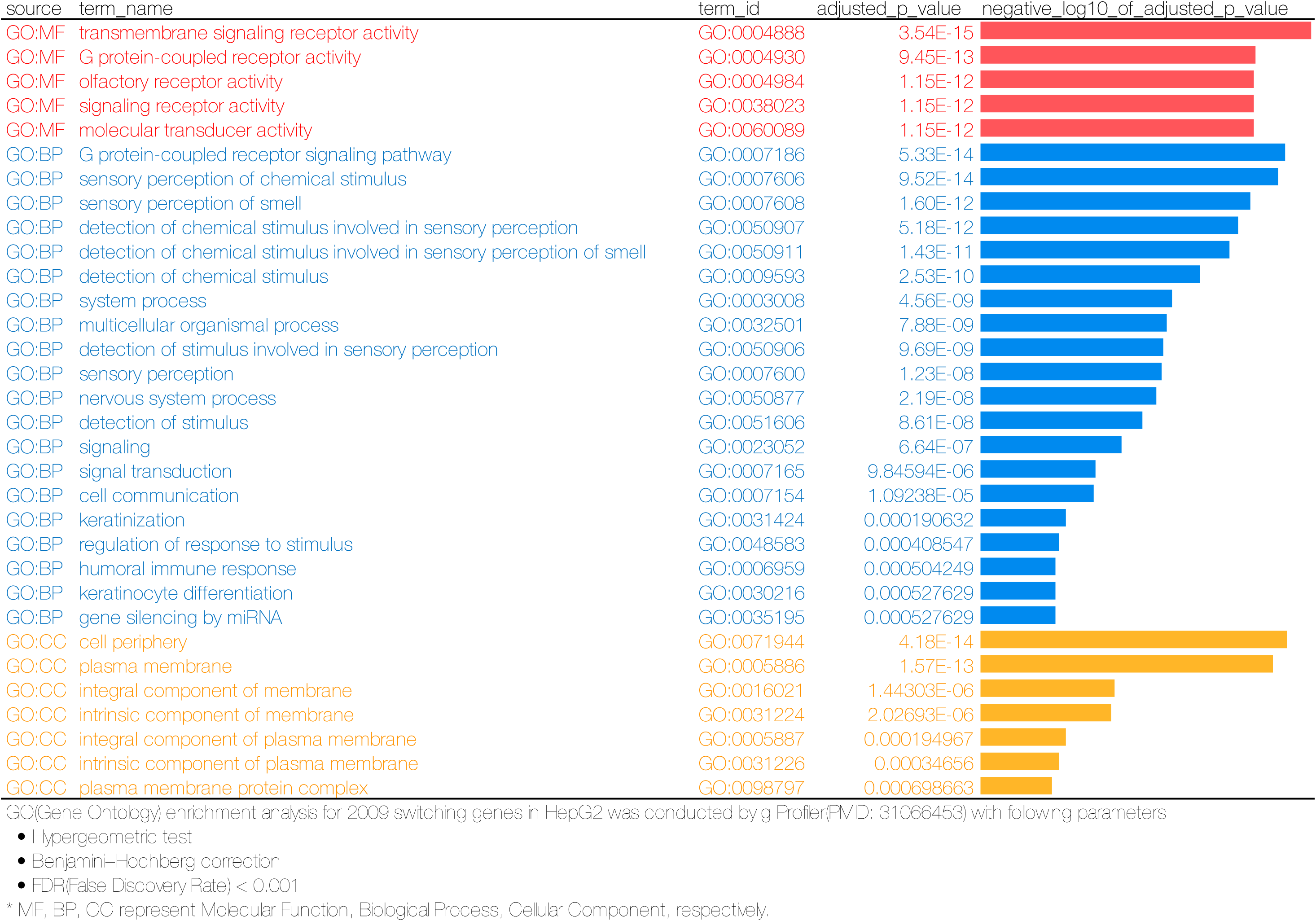
Gene Ontology enrichment analysis for switch genes (HepG2, continued)

**Supplementary Data 5.**
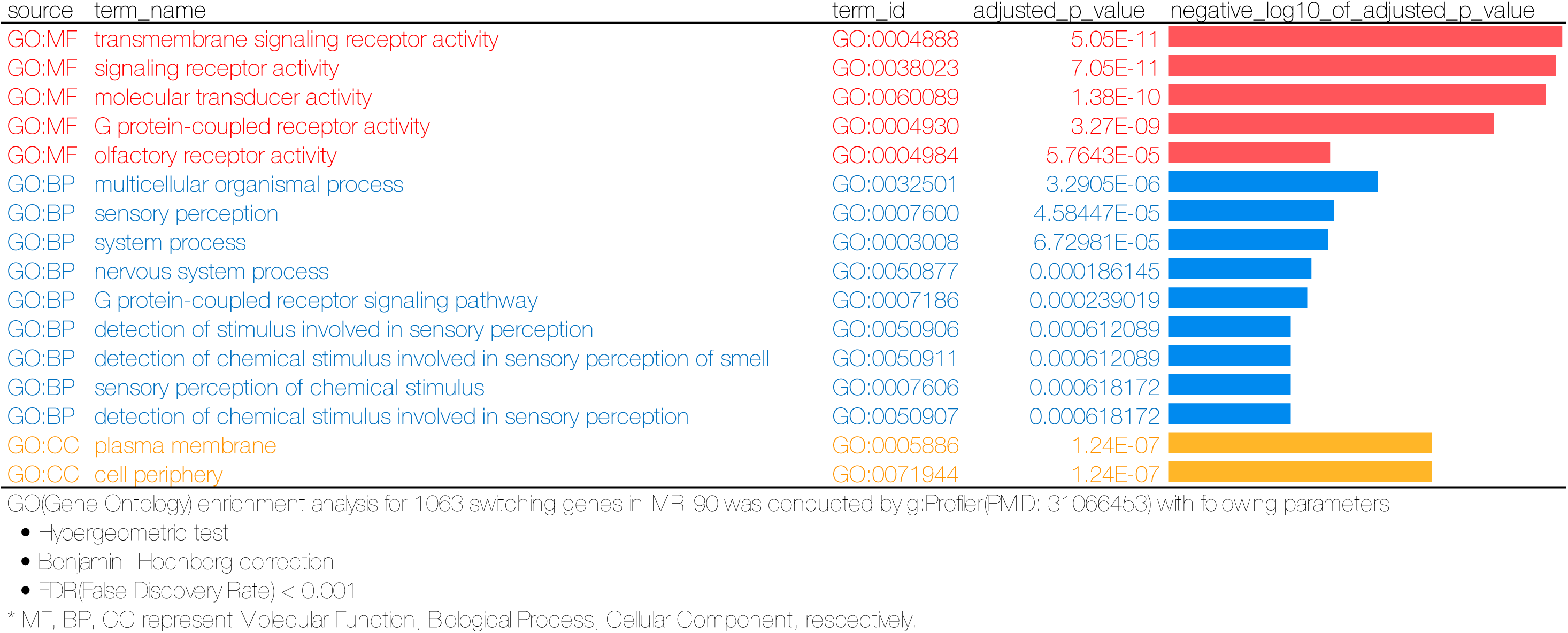
Gene Ontology enrichment analysis for switch genes (IMR-90, continued)

**Supplementary Data 5.**
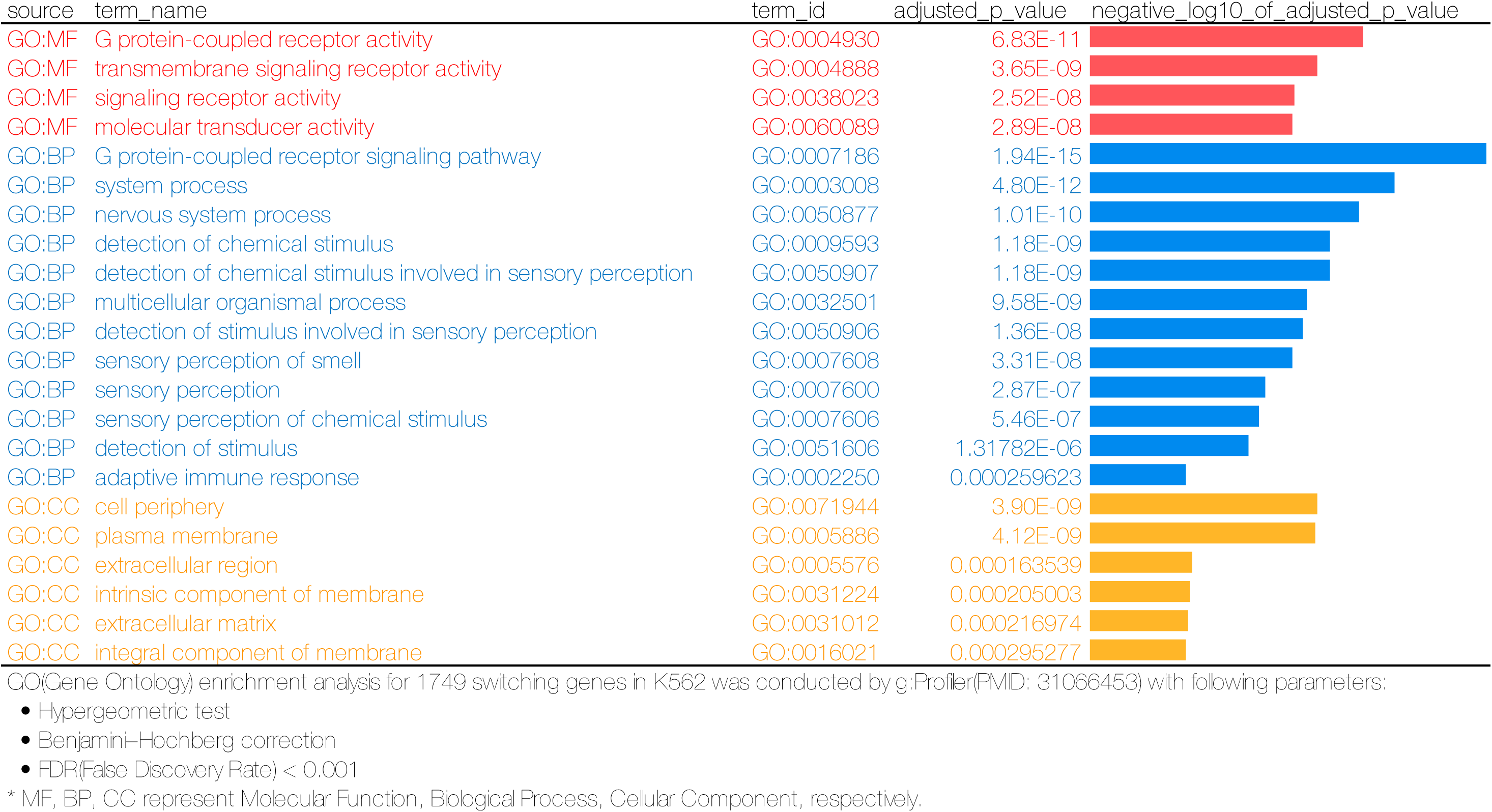
Gene Ontology enrichment analysis for switch genes (K562, continued)

**Supplementary Data 5.**
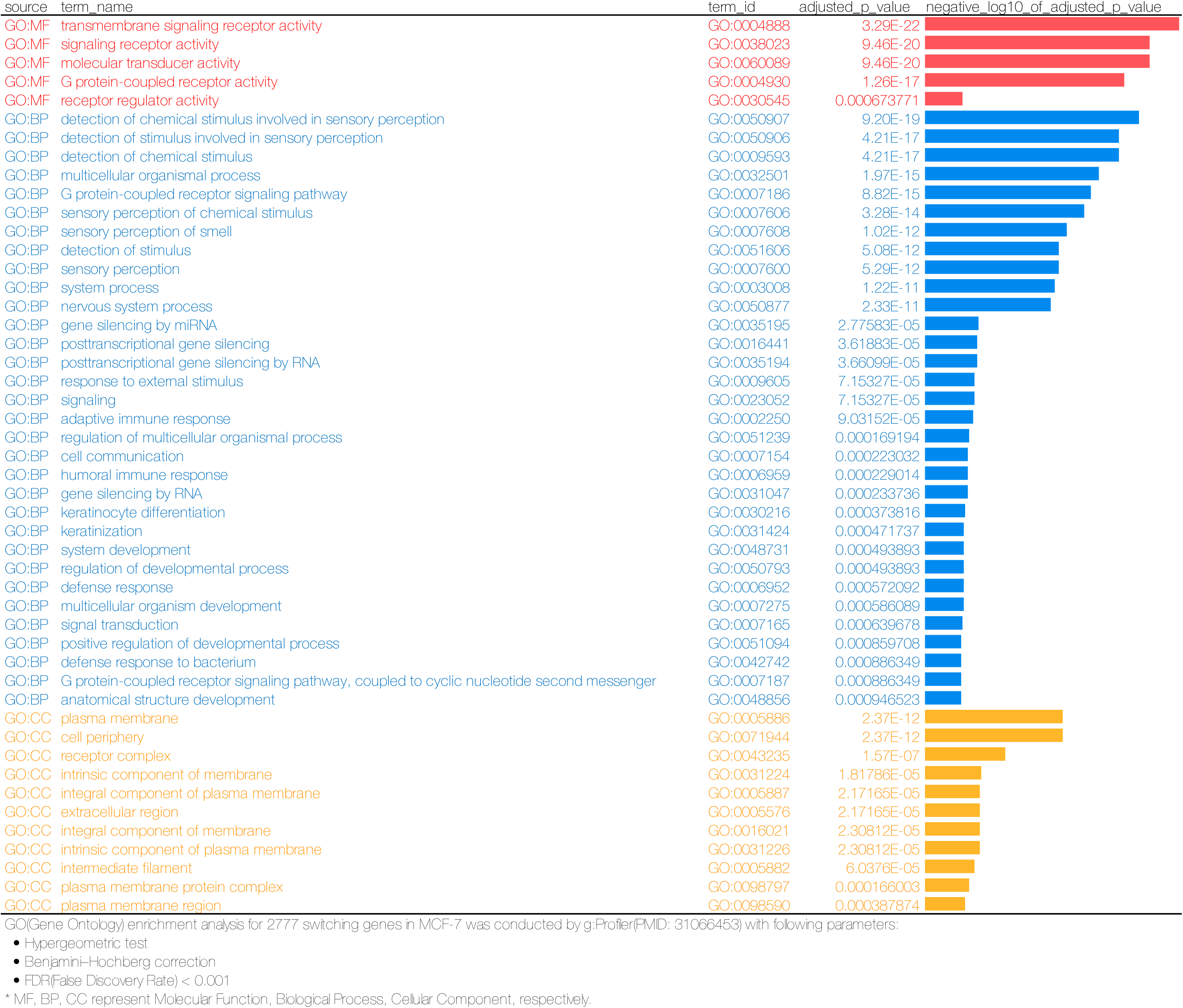
Gene Ontology enrichment analysis for switch genes (MCF-7, continued)

**Supplementary Data 5.**
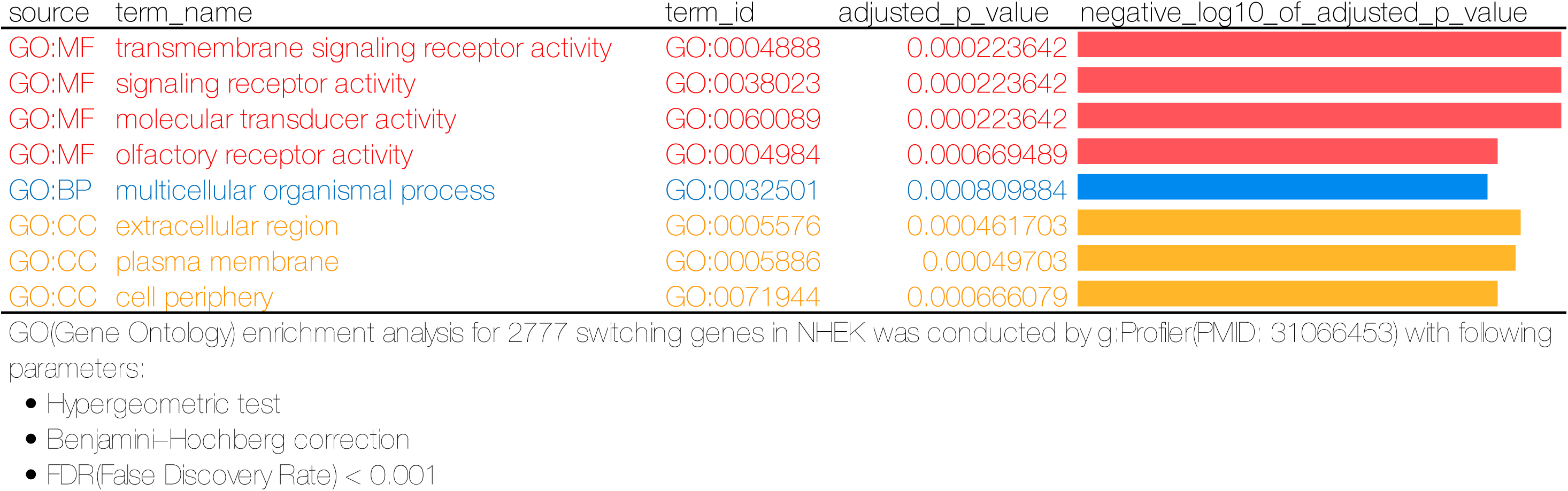
Gene Ontology enrichment analysis for switch genes (NHEK, continued)

**Supplementary Data 5.**
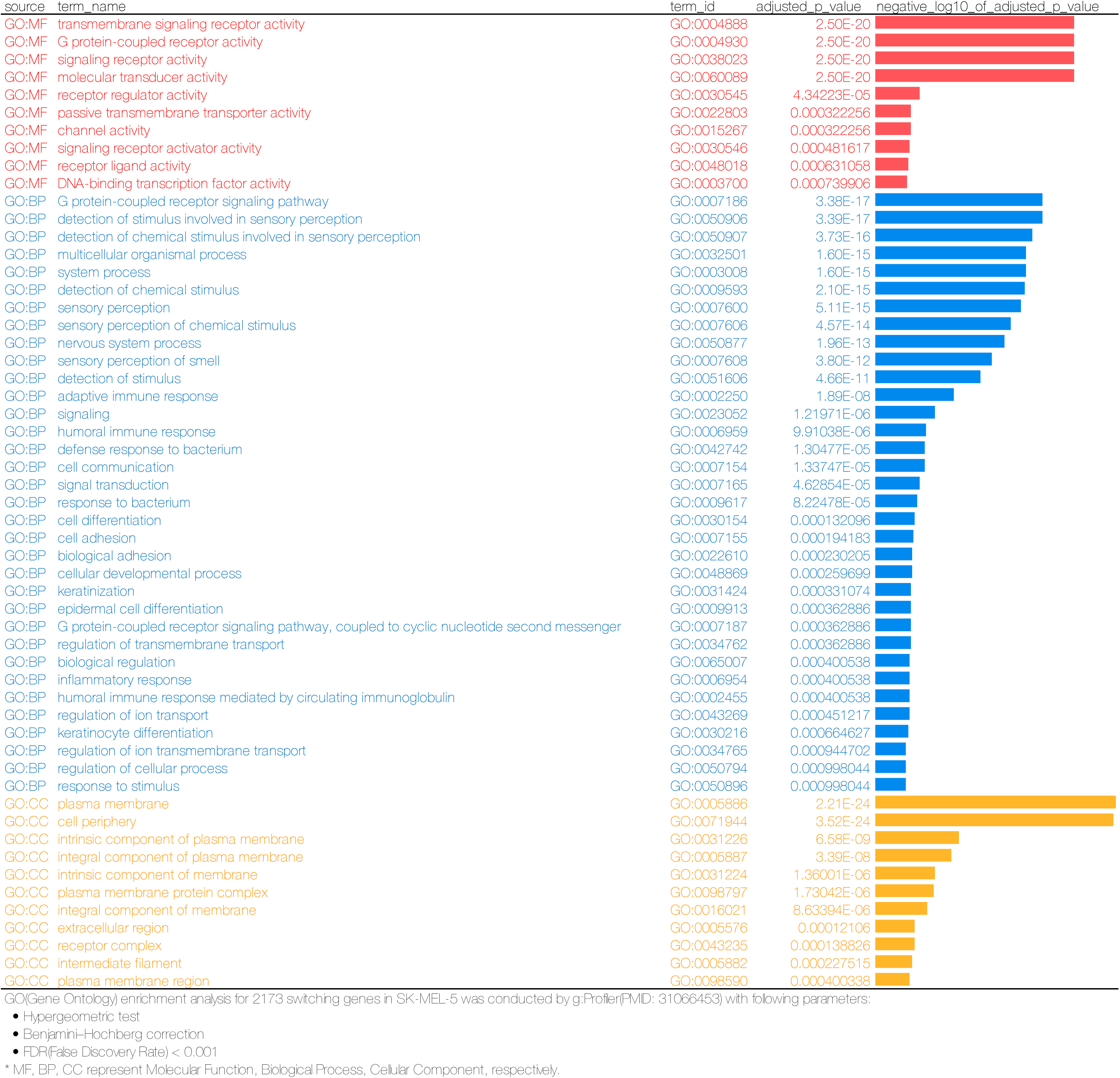
Gene Ontology enrichment analysis for switch genes (SK-MEL-5, continued)

**Supplementary Data 5.**
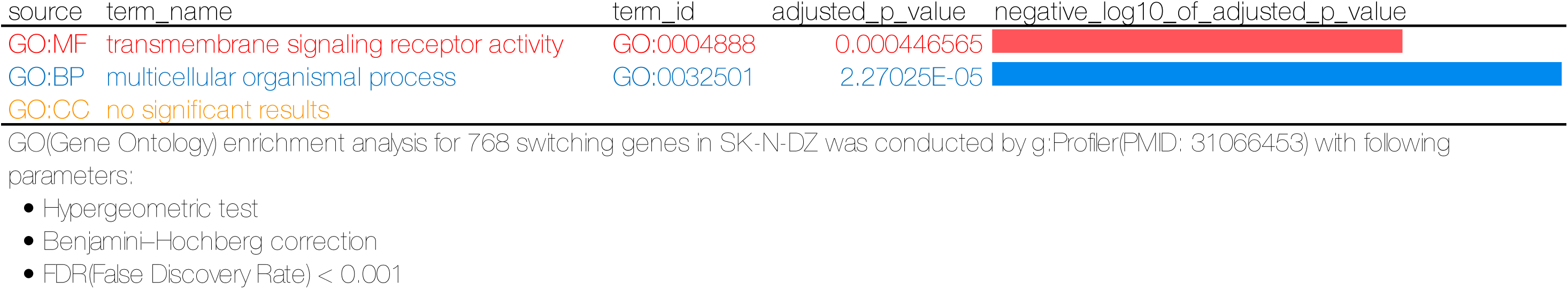
Gene Ontology enrichment analysis for switch genes (SK-N-DZ, continued)

**Supplementary Data 5.**
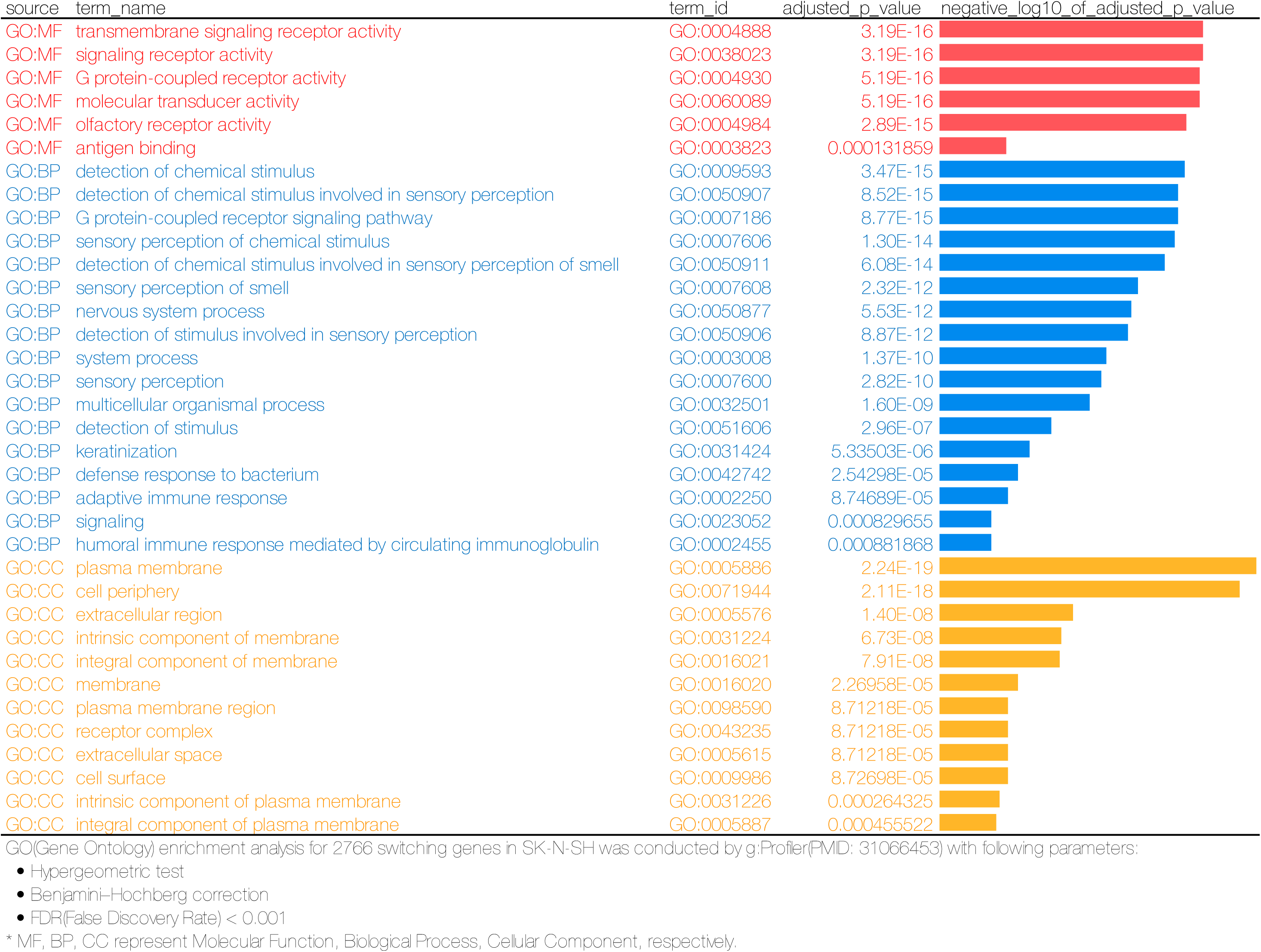
Gene Ontology enrichment analysis for switch genes (SK-N-SH)

**Supplementary Data 6.**
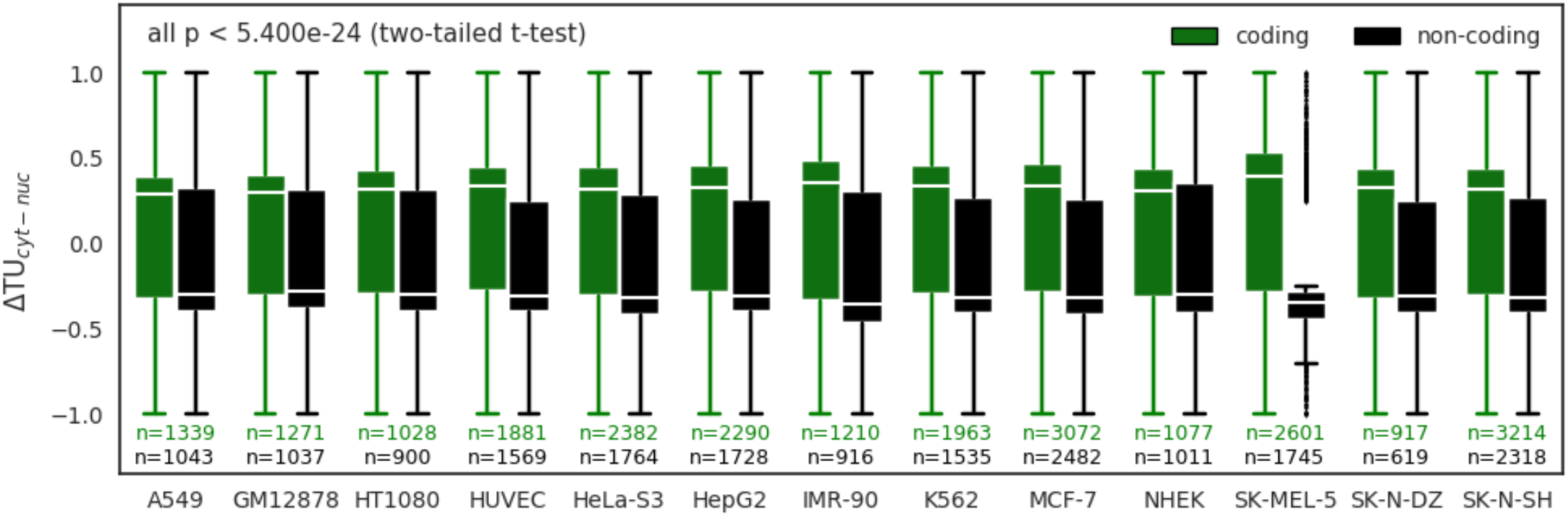
Comparison of ΔTU between protein-coding transcripts (green) and non-coding transcripts (black). Protein-coding transcripts possess higher ΔTU scores, which is consistent with our understanding that the transcript encoding protein is more prone to be located in the cytoplasm. Based on two-tailed t-test, we calculated the significant level of difference between ΔTU in protein-coding and non-coding transcripts in each cell line. The largest p value is 5.400e-24.

**Supplementary Data 7.**
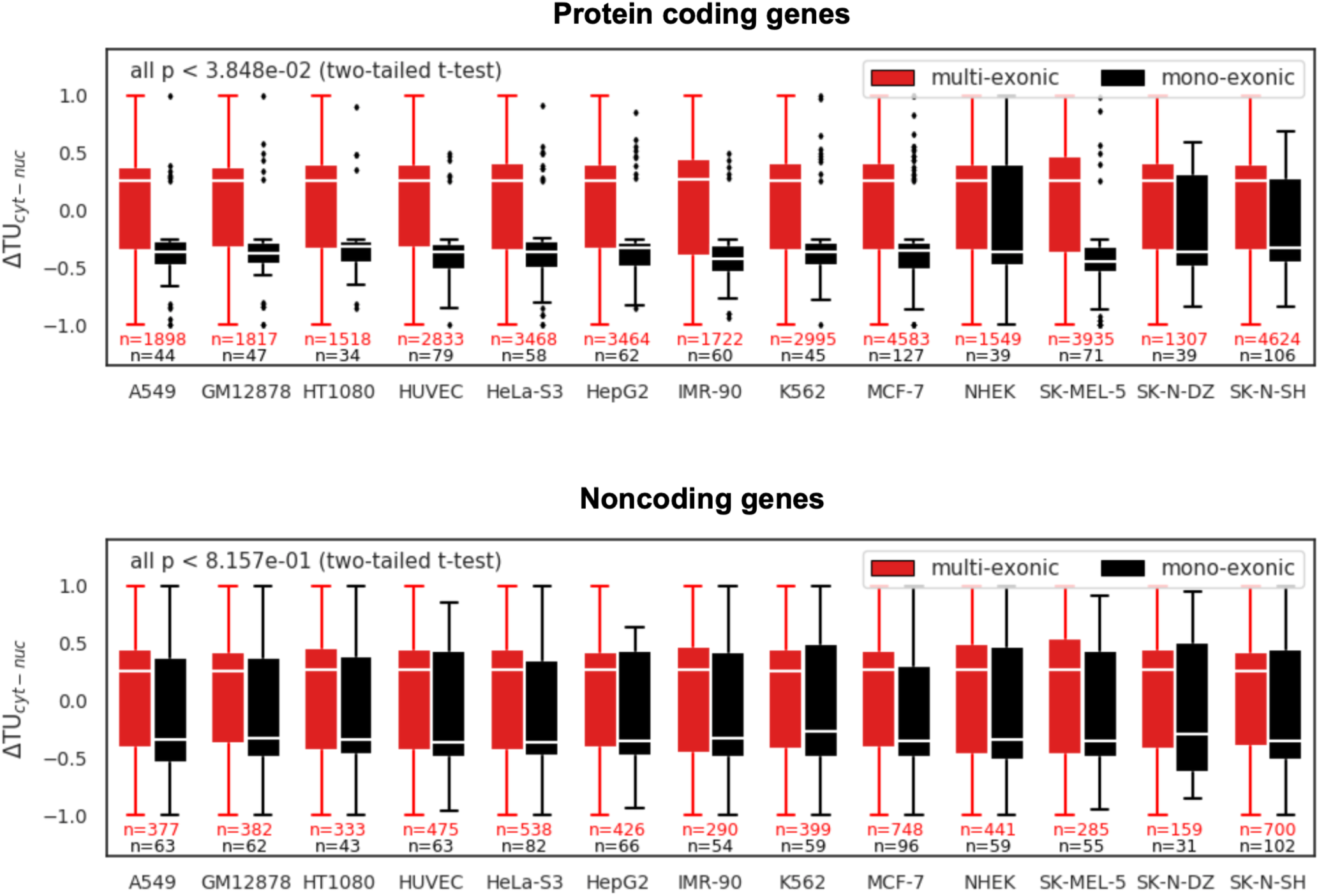
Comparison of ΔTU between mono-exonic (black) and multi-exonic (red) transcripts across 13 cell lines. Top and bottom panels for protein-coding and noncoding genes, respectively. ΔTU shows a significant positive correlation with splicing, indicating that splicing appears to be a dominant factor for RNA export from the nucleus. Based on two-tailed t-test, we calculated the significant level of difference between ΔTU in multi-exonic and mono-exonic transcripts in each cell line. The largest p value is 3.848e-02 and 8.157e-01 for protein-coding and non-coding genes, respectively.

**Supplementary Data 8.**
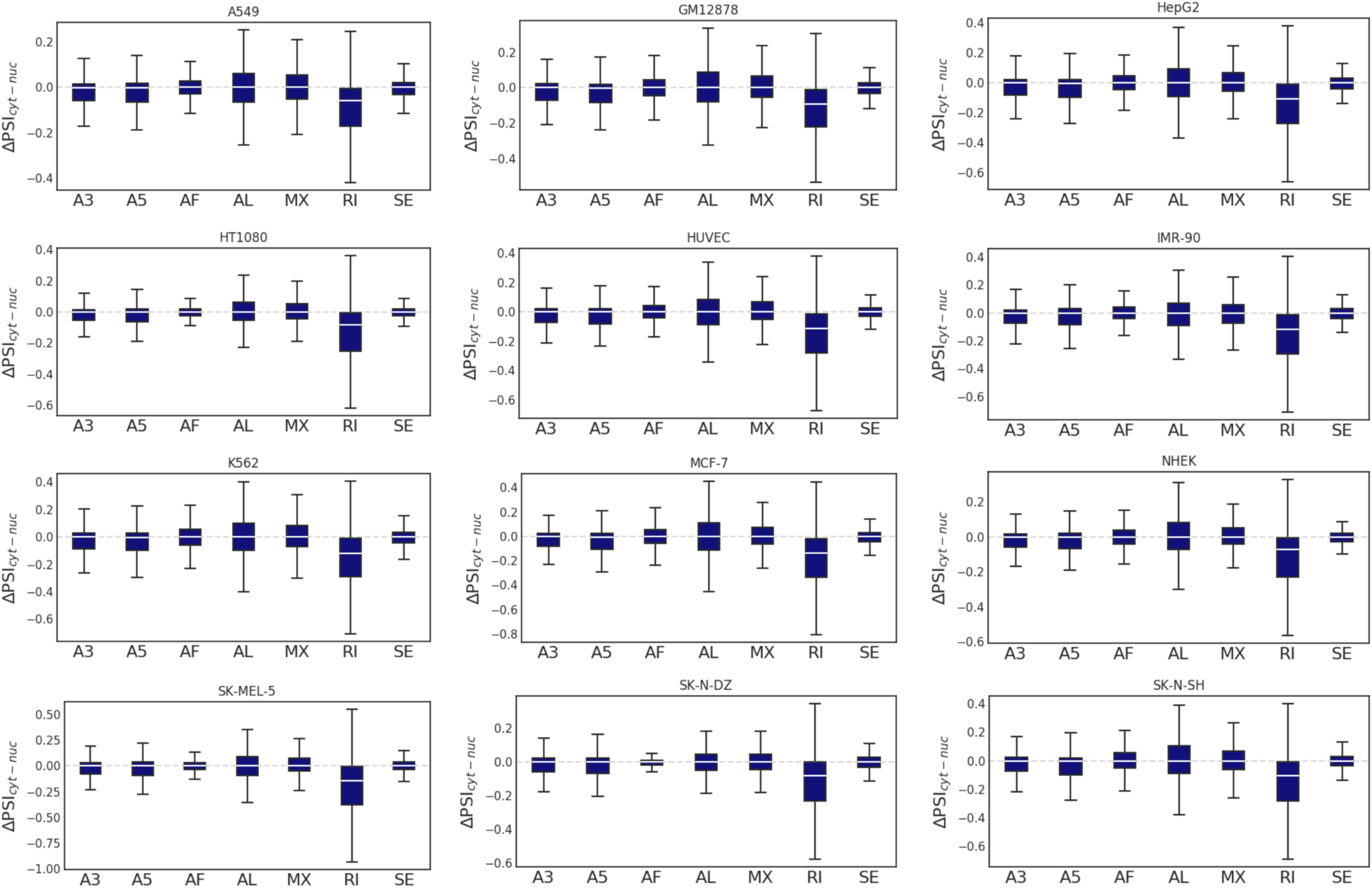
Comparison of alternative splicing patterns between cytoplasmic and nuclear transcripts. (all genes)

**Supplementary Data 9.**
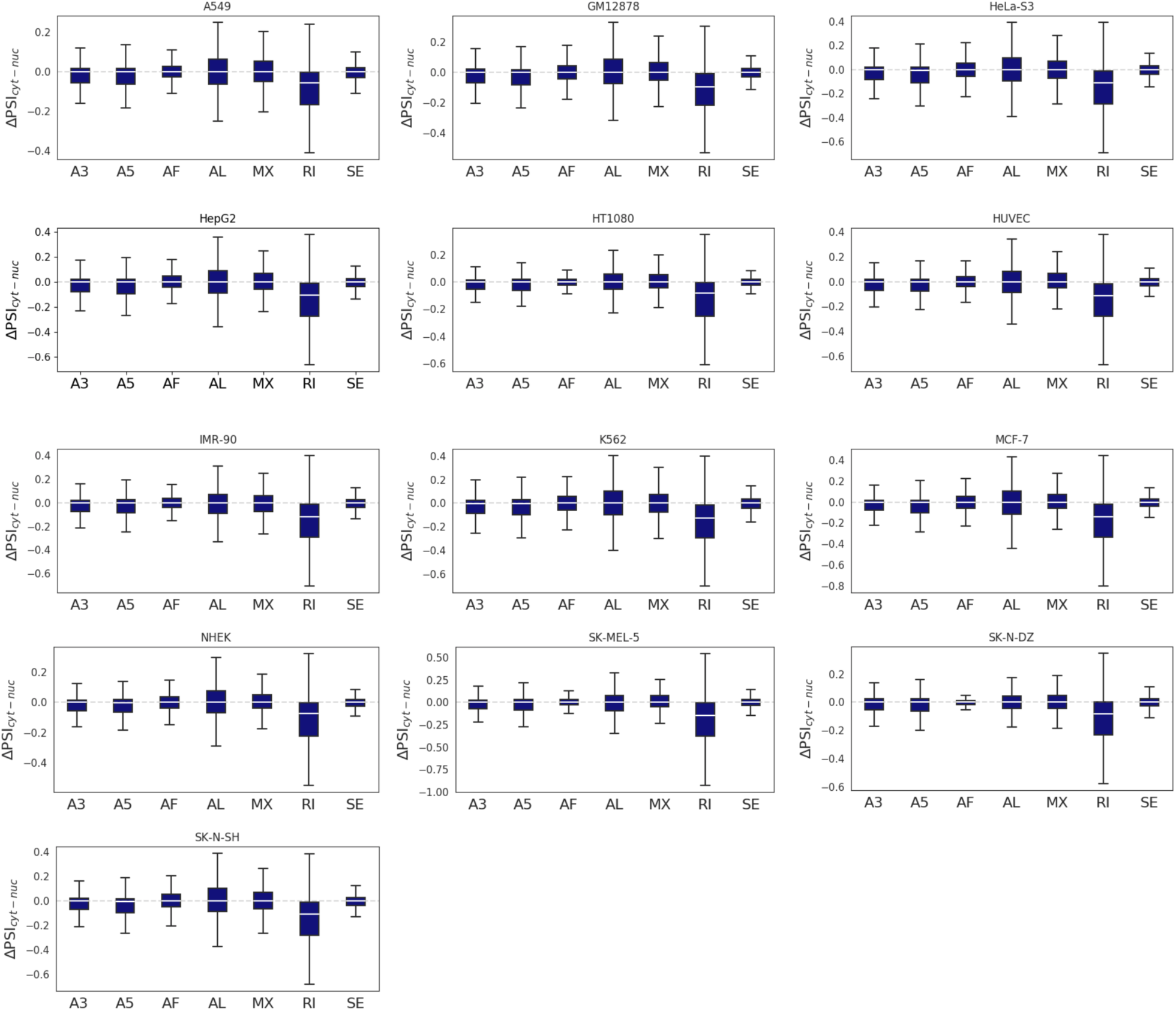
Comparison of alternative splicing patterns between cytoplasmic and nuclear transcripts. (protein-coding genes, continued)

**Supplementary Data 9.**
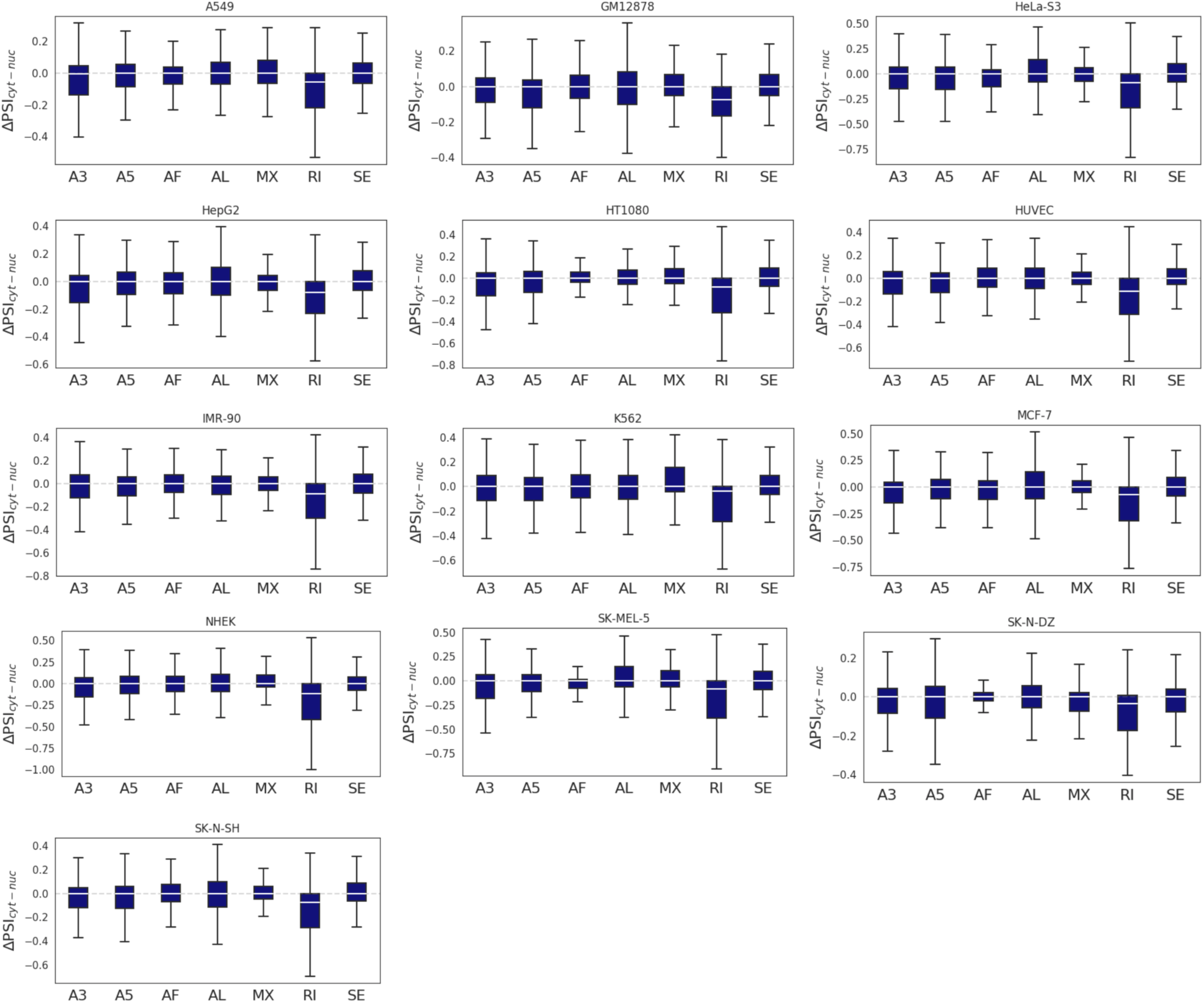
Comparison of alternative splicing patterns between cytoplasmic and nuclear transcripts. (noncoding genes)

**Supplementary Data 10.**
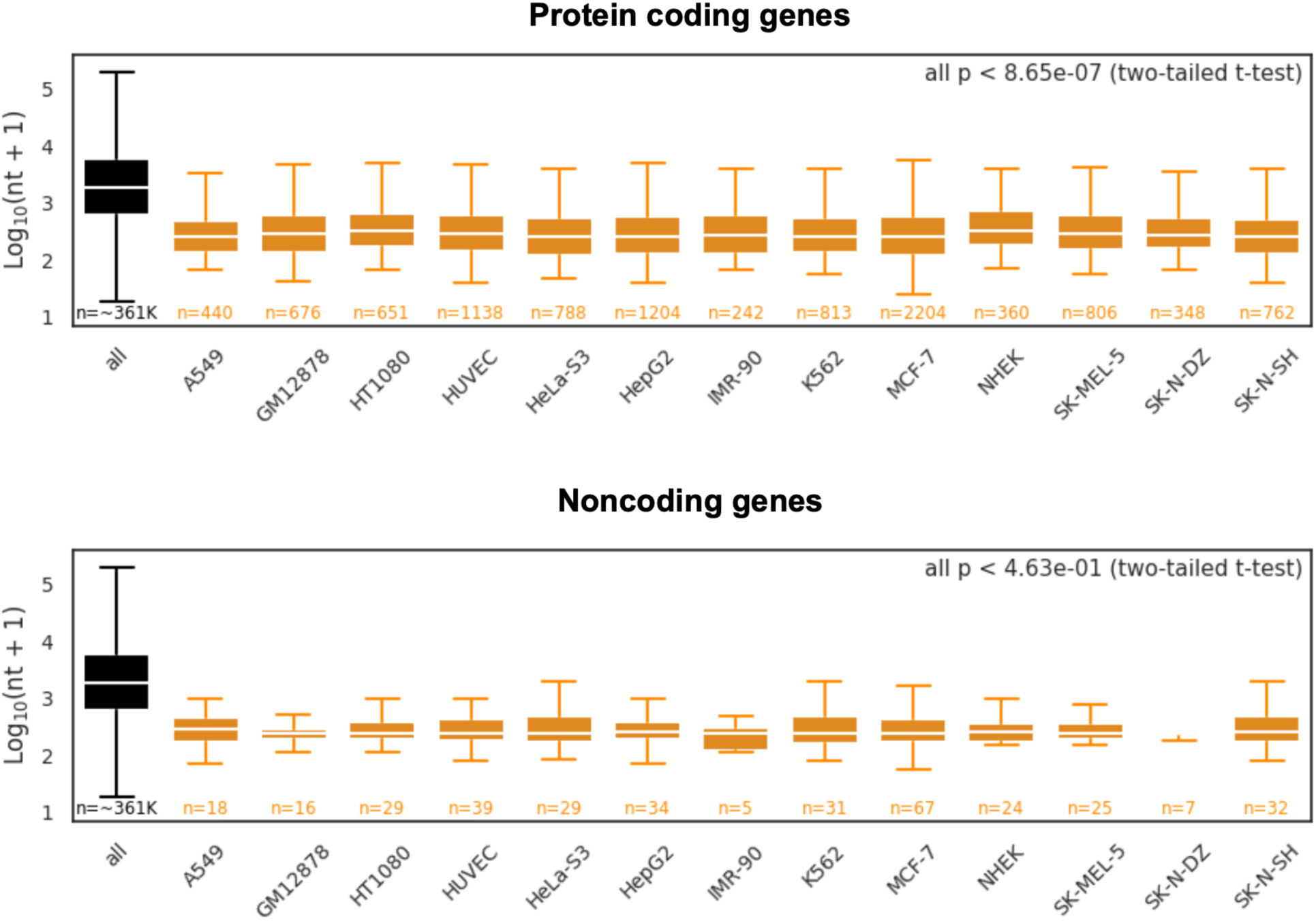
Comparison of length between all introns and nuclear RIs (ΔΨ<0 and p<0.05). Top and bottom panels for protein-coding and noncoding genes, respectively. P values: two-tailed t-test between all introns and nuclear RIs. The largest p value is 8.65e-07 and 4.63e-01 for protein-coding and non-coding genes, respectively.

**Supplementary Data 11.**
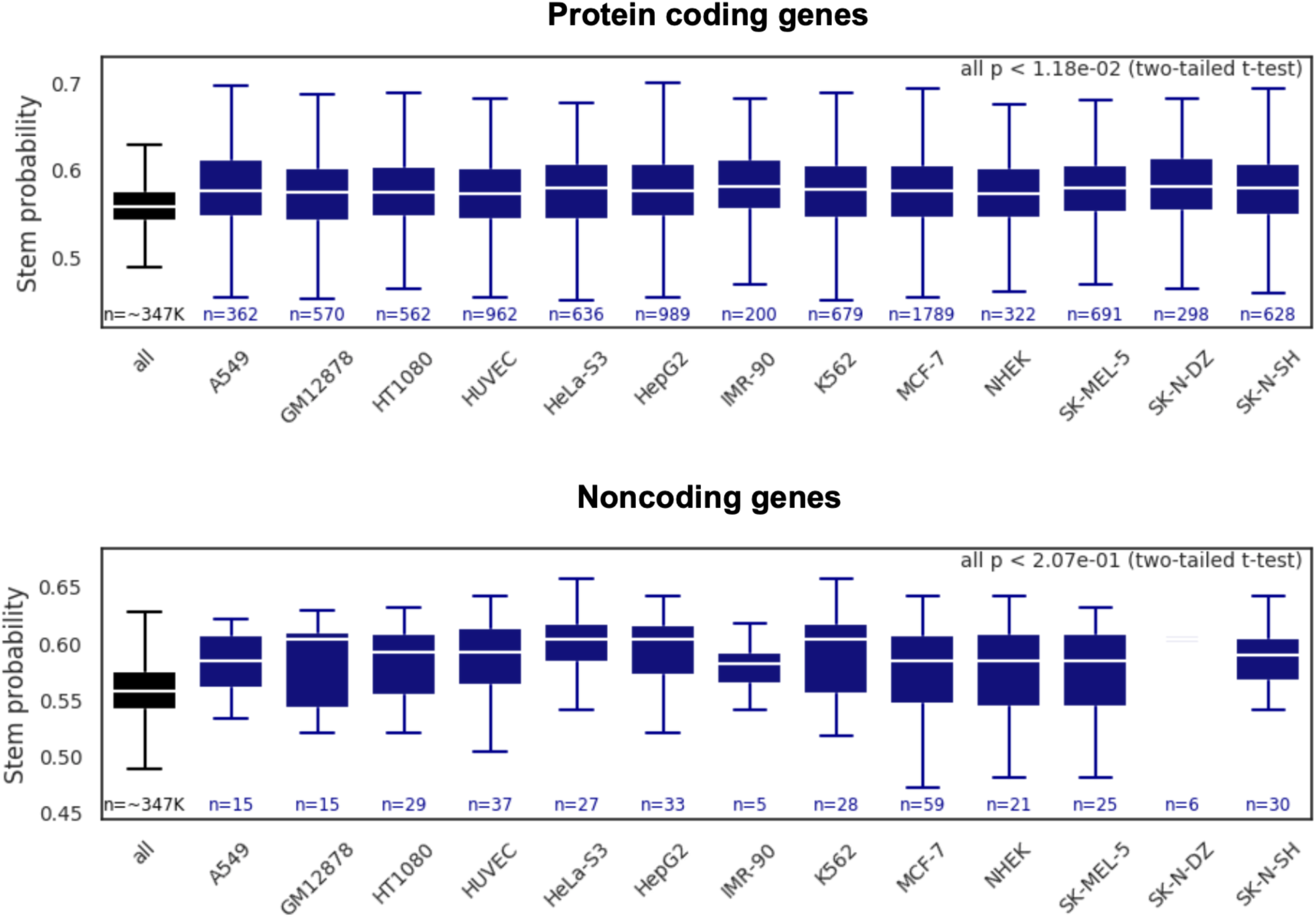
Comparison of RNA secondary structure (stem probability) between all introns and nuclear RIs (ΔΨ<0 and p<0.05). Top and bottom panels for protein-coding and noncoding genes, respectively. P values: two-tailed t-test between all introns and nuclear RIs. The largest p value is 1.18e-02 and 2.07e-01 for protein-coding and non-coding genes, respectively.

